# Dihydroxyhexanoic acid biosynthesis controls turgor in pathogenic fungi

**DOI:** 10.1101/2024.08.07.606736

**Authors:** Naoyoshi Kumakura, Takayuki Motoyama, Keisuke Miyazawa, Toshihiko Nogawa, Katsuma Yonehara, Kaori Sakai, Nobuaki Ishihama, Kaisei Matsumori, Pamela Gan, Hiroyuki Koshino, Takeshi Fukuma, Richard J. O’Connell, Ken Shirasu

## Abstract

Many plant pathogenic fungi penetrate host surfaces mechanically, using turgor pressure generated by appressoria, specialized infection cells. These appressoria develop semipermeable cell walls and accumulate osmolytes internally to create turgor by osmosis. While melanin is known to be important for turgor generation, the mechanism for wall semipermeability has remained unclear. Here we identify *PKS2* and *PBG13*, by reverse genetics, as crucial for forming the semipermeable barrier in anthracnose and rice blast fungi. These genes encode enzymes that synthesize 3,5-dihydroxyhexanoic acid polymers essential for the cell wall properties. Deleting these enzymes impairs cell wall porosity, abolishing turgor and pathogenicity without affecting melanization. Our findings uncover a novel mechanism of turgor generation, linking enzyme function to pathogen penetration and disease potential, presenting new targets for disease control.

## Main Text

Fungal pathogens, such as *Colletotrichum* species and *Magnaporthe oryzae* (*Mo*), are devastating agricultural pests (*1*, *2*). *Colletotrichum* fungi, with approximately 260 reported species, damage an enormous range of crops including major cereals, fruits, and forage crops (*3*). *Mo* causes annual losses of 10-30% in global rice production and has recently caused serious epidemics in wheat (*4*). Understanding the pathogenic mechanisms of these fungi is an urgent challenge for food security. These pathogens use specialized cells known as appressoria (*5*), which develop high turgor pressure to breach the plant surface, thereby enabling infection (*6–9*). The spores of these fungi adhere to the plant surface, where they germinate to form appressoria, which are darkly pigmented due to the accumulation of melanin, a biopolymer derived from 1,8-dihydroxynaphthalane (DHN), in their cell walls (*10–13*). Appressoria accumulate osmolytes internally (*14*), leading to high osmotic pressure and subsequent water absorption, thereby generating high turgor pressure through hydrostatic forces (∼8 MPa) (*15*). This process suggests that the appressorial cell wall serves as a semipermeable barrier, allowing water permeation while restricting osmolyte passage (*16*). Although melanin was shown to play a crucial role in appressorial turgor generation in the rice blast fungus, its presence in hyphal cells that do not generate high turgor (*15*, *17*) suggests that other specific factors contribute to forming the semipermeable barrier. Consistent with this, appressoria of a *Colletotrichum* species can generate turgor pressure even in the absence of melanin (*18*), suggesting additional unknown factors involved in this mechanism. Here, we report a novel polyketide that confers semipermeability to the fungal cell wall and the two enzymes required for its biosynthesis.

### *PKS2* and *PBG13* are essential for infection

To identify the factors crucial for appressorial functionality, we employed a reverse genetic approach using *Colletotrichum orbiculare* (*Co*), a transformable pathogen causing anthracnose disease in cucurbits (*19*, *20*). Our previous study highlighted that many genes encoding secondary metabolite (SM) synthesis enzymes and secreted proteins are upregulated during infection (*21*). Here, we focused on these SM biosynthetic enzymes of *Co*, which remain largely unexplored, except for *CoPKS1*, a polyketide synthase involved in DHN-melanin synthesis (*22*). We identified 73 <u>s</u>econdary <u>m</u>etabolism <u>k</u>ey enzyme-encoding <u>g</u>enes (SMKGs) in the *Co* genome (*23*), including those encoding polyketide synthases (PKSs) and dimethylallyltryptophan synthase (DMATS) among others (fig. S1A, table S1). Using RNA-seq analysis (*24*), we identified 23 SMKGs that exhibit specific upregulation during infection (Fig. 1A, fig. S1B). Four SMKGs (*CoDMATS4*, *CoNRPSL9*, *CoNRPS8*, and *CoPKS2*) are selected due to their high expression levels at 1 day-post-inoculation (DPI), when appressoria are forming. Targeted knock-out of these genes using CRISPR-Cas9 (*25*, *26*) revealed that *CoPKS2*, encoding a highly-reducing polyketide synthase, plays a pathogenicity-specific role (Fig. 1B-C, fig. S1C-F). *Copks2* was unable to cause lesions, but reintroduction of the *CoPKS2* gene recovered the wild-type (WT) disease phenotype. Additionally, no differences were detected in colony or spore morphology between *Copks2* and the parental strain, *Co*0051 (*27*), the latter retaining similar pathogenicity to WT, indicating that *CoPKS2* is specifically required for infection (fig. S1G-J).

**Fig. 1.**
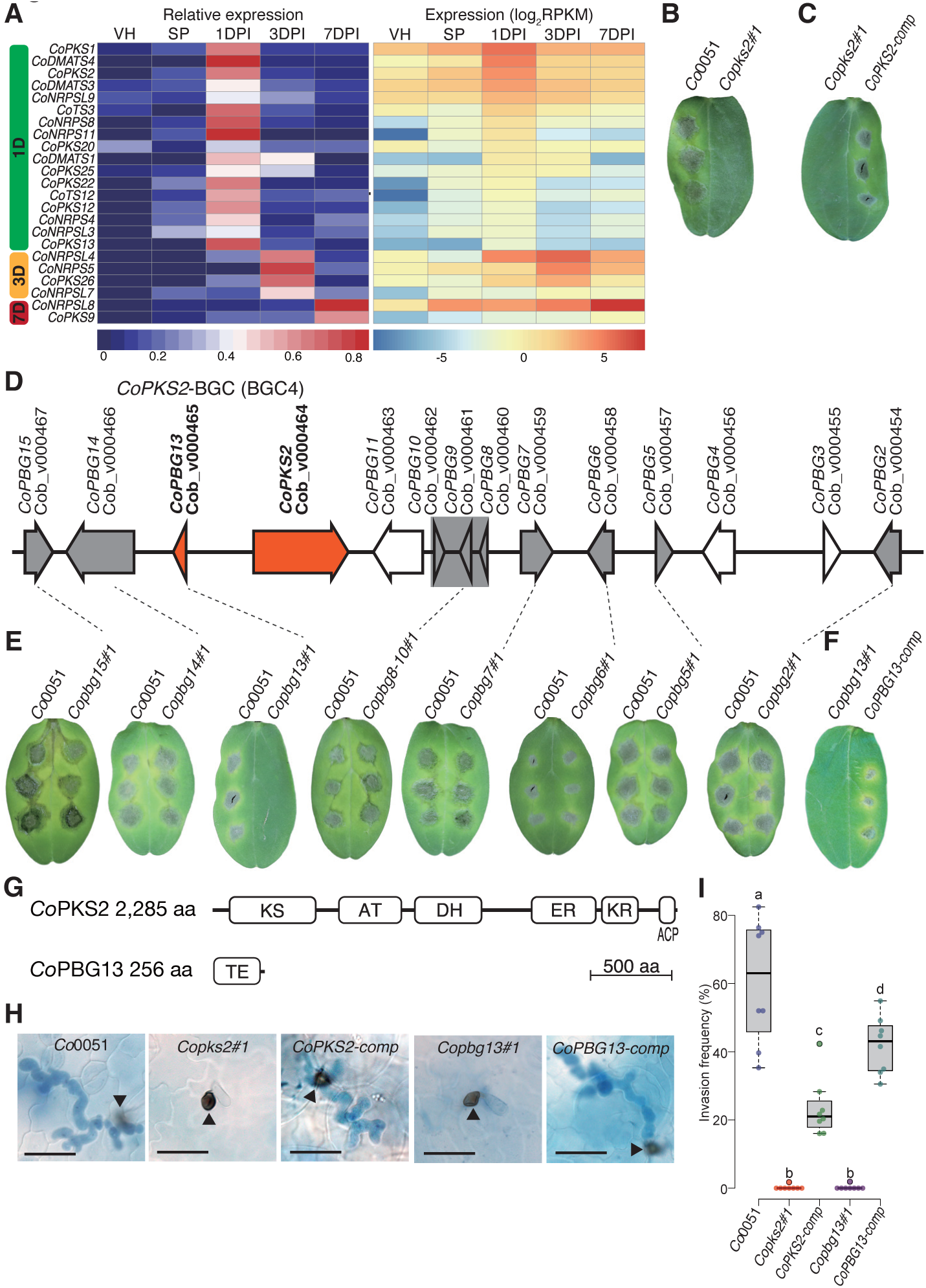
*Co*PKS2 and *Co*PBG13 enzymes are required for *Co* infection. (**A**) Twenty-three *Co*SMKGs are up-regulated during infection. Vegetative hyphae (VH) and spore (SP) were non-infection controls. (**B**) *Copks2* mutants showed no disease symptoms on cucumber leaves. Four leaves were used for each strain. (**C**) Re-introduction of the *CoPKS2* gene restored the disease phenotype in *Copks2*. (**D**) Schematic of the *CoPKS2*-BGC. Grey and orange regions/genes were knocked out in (E). (**E**) *Copbg13* mutants exhibited the same deficiency in disease symptom production as *Copks2*. (**F**) Re-introduction of the *CoPBG13* gene restored the disease phenotype in *Copbg13*. (**G**) Predicted domains of *Co*PKS2 and *Co*PBG13. KS: ketosynthase, AT: acyltransferase, DH: dehydratase, ER: enoyl reductase, KR: ketoreductase, ACP: acyl carrier protein, TE: thioesterase, aa: amino acids. (**H**) Invasive hyphae (stained blue) emerging from appressoria were not observed in *Copks2* and *Copbg13*. Black triangles indicate appressoria. Scale bars represent 20 µm. (**I**) Invasion frequency was calculated as the percentage of appressoria that formed invasive hyphae. Each datapoint was calculated from at least 50 appressoria. Eight inoculation sites were used for each strain (n=8). Different characters indicate a statistically significant difference (p<0.05, Tukey HSD). In (B-C, E-F, H-I), three independent experiments yielded similar results.

Fungal SMKGs are typically clustered in specific genomic regions known as biosynthetic gene clusters (BGCs) (*28*). In silico analysis predicted 58 potential BGCs in the *Co* genome, with *CoPKS2* located within BGC4 comprising 23 genes named *Co<u>P</u>KS2-<u>B</u>GC <u>g</u>enes* (*CoPBGs*) (Fig. 1D, Data S1). This led us to speculate that other BGC4 genes in addition to *CoPKS2* are involved in the biosynthesis of an SM. To test this, ten genes in BGC4 showing a similar expression pattern to *CoPKS2* were knocked-out, and the impact on pathogenicity was assessed (Fig. 1E-F, fig. S2A-D). Notably, only mutants lacking *CoPBG13* demonstrated no pathogenicity, which was rescued by introducing the *CoPBG13* genomic sequence (*CoPBG13-comp*). No differences in colony or spore morphology were observed between *Copbg13* and *Co*0051, similar to *Copks2* (fig. S1G-J). While *Co*PKS2 lacks the thioesterase domain, typically involved in cleaving polyketide products from PKS, this domain was present in *Co*PBG13 (Fig. 1G), suggesting that *Co*PBG13 may be crucial for releasing the SM synthesized by *Co*PKS2.

Appressoria-mediated penetration of the cucumber epidermis allows *Co* to form invasive hyphae within the leaf. In strains containing *CoPKS2* and *CoPBG13* (*Co*0051, *CoPKS2-comp,* and *CoPBG13-comp*), appressoria-derived invasive hyphae were observed, while no such structures were detected in strains lacking these genes (*Copks2* and *Copbg13*) (Fig. 1H-I). No significant differences were noted in the morphology, melanization, and frequency of appressoria formation among these strains (fig. S2E-F). These findings emphasize the essential role of *CoPKS2* and *CoPBG13* in appressorial function.

### Conservation of *PKS2* and *PBG13*

Next, we investigated conservation of the *PKS2* and *PBG13* gene pair across a range of fungal phytopathogens, including multiple *Colletotrichum* species and *Mo* known for their melanized appressoria. The gene pair was conserved in all eight examined *Colletotrichum* species, each possessing one pair (Fig. 2A, fig. S3, table S2). *Mo* has two pairs, and other plant pathogens, including *Verticillium dahliae* and *Botrytis cinerea*, each have one pair. To explore the functional conservation of *CoPKS2* and *CoPBG13* in other *Colletotrichum* species, we focused on *C. higginsianum* (*Ch*) (*29*), a transformable pathogen of Brassicaceae plants including *Arabidopsis thaliana*. Using a CRISPR-Cas9-based approach (*30*) (fig. S4A-H), we demonstrated that *Chpks2* and *Chpbg13* lost pathogenicity (Fig. 2B-C), without showing any differences in colony or spore morphology (fig. S4I-L). Notably, pathogenicity in *Copks2* was restored by knocking-in *ChPKS2* (Fig. 2D), indicating the functional equivalence between *ChPKS2* and *CoPKS2*. Similarly, *Mopks2a* and *Mopbg13a* mutants lost pathogenicity on rice, while *Mopks2b* showed no such phenotype (Fig. 2E-F, fig. S5A-B). Pathogenicity was restored in the respective complemented strains. Moreover, *CoPKS2* introduction restored pathogenicity in *Mopks2a*, confirming the functional similarity between *Mo*PKS2A and *Co*PKS2 (Fig. 2G). These cross-complementation results collectively demonstrated functional conservation of *PKS2* and *PBG13* across three fungal species.

**Fig. 2.**
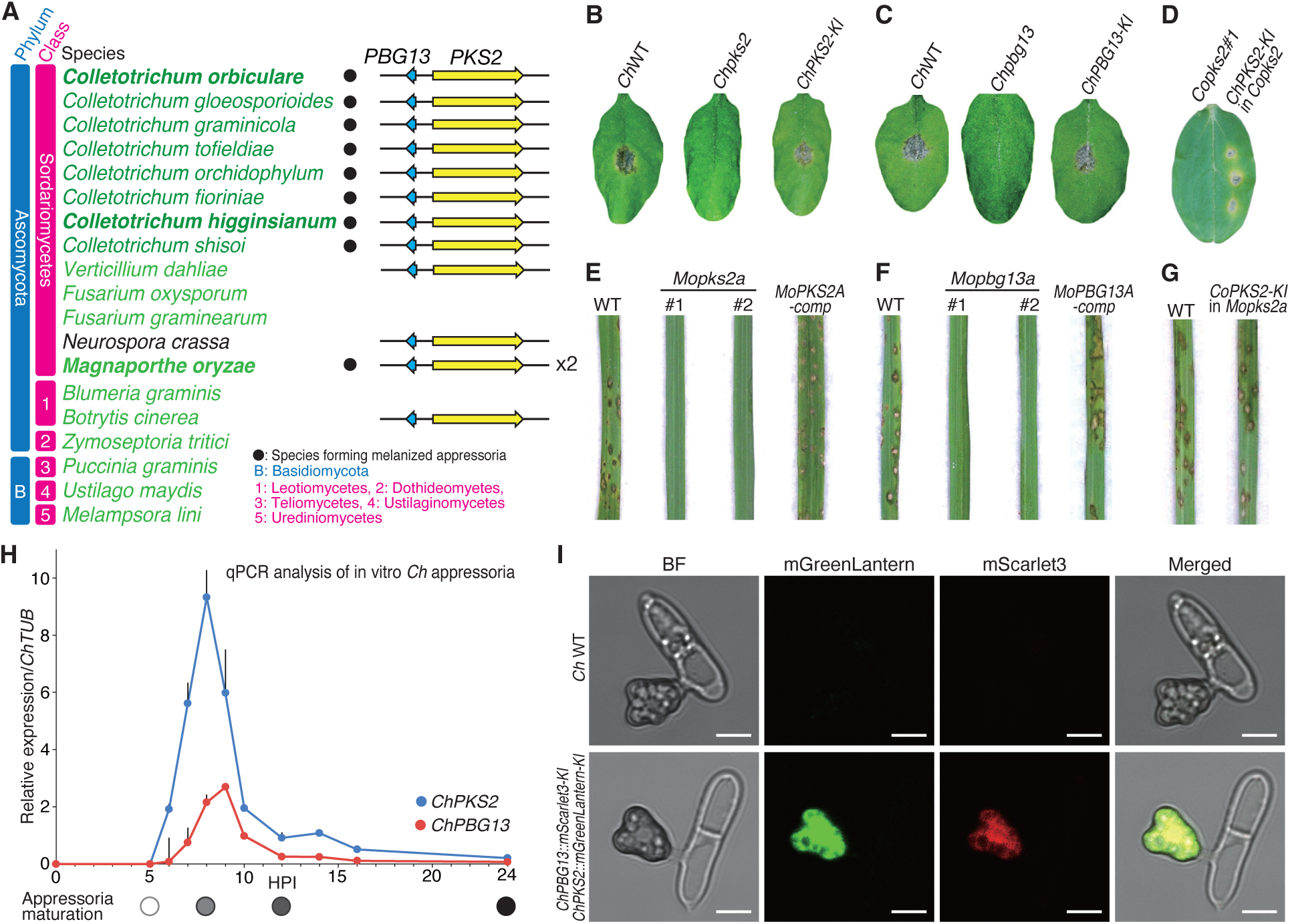
Homologs of *Co*PKS2 and *Co*PBG13 are essential for the virulence of *Ch* and *Mo*. (**A**) Conservation of syntenic *PKS2* and *PBG13* genes. “x2” indicates two sets of *PKS2-PBG13* genes were detected. (**B**) A *Chpks2* mutant showed a loss of virulence phenotype. *Ch* was inoculated onto *A. thaliana* Col-0 leaves. Four leaves were used for each strain. (**C**) A *Chpbg13* mutant showed a loss of virulence phenotype. Experimental conditions were described in (B). (**D**) Complementation of *Copks2* by *ChPKS2*. Five cucumber cotyledons were used. (**E**) *Mopks2a* mutants showed a loss of virulence phenotype. The *Mo* Kita-1 strain was sprayed onto 10-14 leaves of the rice cultivar Nipponbare. (**F**) *Mopbg13a* mutants showed a loss of virulence phenotype. (**G**) *CoPKS2* gene complemented the *Mopks2a* loss of virulence phenotype. In F and G, experiments were performed in the same way as described in (E). (**H**) *ChPKS2* and *ChPBG13* are up-regulated during the early development of appressoria. Transcript levels of *Ch*WT appressoria were quantified with qPCR (n=3), with the *ChTUB* as a reference. Error bars: +SE. (**I**) *Ch*PKS2-mGreenLantern and *Ch*PBG13-mScarlet3 localize to the appressorial cytosol. Appressoria were observed at 8 HPI. Scale bars: 5 µm. In (B-G), three independent experiments yielded similar results.

### PKS2 and PBG13 form a semipermeable barrier

To understand the roles of *PKS2* and *PBG13* in appressorium development, we assessed the upregulation of these genes in *Ch,* a system that allows synchronized mass-production of appressoria (*30*, *31*). The development of the appressorium occurs within a day (*32*), beginning with germ-tube emergence from the spore on artificial substrata or plant tissue (∼4 hours post inoculation [HPI]), followed by formation of a transparent appressorium from the germ tube (∼6 HPI), and culminating in melanization and pressurization (∼24 HPI). Interestingly, expression of *ChPKS2* and *ChPBG13* peaked at 8 HPI and then sharply declined, suggesting their involvement at an early stage of appressorium development (Fig. 2H). Consistently, we detected *Ch*PKS2 and *Ch*PBG13 fused with fluorescent proteins (mGreenLantern (*33*) and mScarlet3 (*34*)) at 8 HPI (Fig. 2I). Importantly, this transgenic strain retained pathogenicity on the host plant, *A. thaliana*, and successful expression of each fluorescent fusion protein was confirmed by immunoblotting (fig. S5C-E). Confocal microscopy observation revealed that both *Ch*PKS2-mGreenLantern and *Ch*PBG13-mScarlet3 localized throughout the cytosol (Fig. 2I). This localization suggests that the metabolite products of *Ch*PKS2 and *Ch*PBG13 are synthesized in the cytoplasm during early appressorium formation. Similar localization patterns of *Co*PKS2-EGFP and *Co*PBG13-mCherry were observed in *Co* appressoria (fig. S5F-M), corroborating the functional conservation across species.

Next, using two different approaches, we investigated whether PKS2 and PBG13 are involved in the production of turgor pressure. We first measured the apparent stiffness (*k*_as_) of the appressoria by atomic force microscopy (AFM), assuming that turgor pressure modulates their mechanical properties (*35*) (fig. S6). The *k*_as_ of *Copks2* and *Copbg13* significantly decreased compared to *Co*0051, while in *CoPKS2-comp* and *CoPBG13-comp*, the *k*_as_ recovered (Fig. 3A-B), indicating *Co*PKS2 and *Co*PBG13 contribute to the stiffness of appressoria, probably by regulating turgor pressure. We then evaluated appressorial turgor pressure with an incipient cytorrhysis assay (*15*, *36*), employing a concentration gradient of a large osmolyte, PEG-6000 (6 kDa), that cannot pass through the appressorial cell wall. The high osmotic pressure induced by PEG-6000 causes the appressoria to either collapse through a process known as cytorrhysis or form air bubbles in the cytoplasm due to water loss, a process termed cavitation (*36*) (fig. S7). Defining appressorial turgor as the osmotic pressure at which 50% of the appressoria undergo cytorrhysis or cavitation (*15*), we found that the appressoria of *Co*0051 and *CoPKS2-comp* exhibited a turgor of approximately 3.7 MPa (Fig. 3C). However, in *Copks2*, even at higher osmotic pressures, less than 50% of the cells showed cytorrhysis or cavitation, indicating *Copks2* lost the ability to generate high turgor. Instead, *Copks2* showed increased plasmolysis, indicating osmolyte permeation through the cell wall (Fig. 3D-E, fig. S7). Similar results were obtained for *Copbg13* and *CoPBG13-comp* (Fig. 3E-G). Our results demonstrate that *Co*PKS2 and *Co*PBG13 contribute to turgor generation, conceivably by reducing the pore size of the cell wall.

**Figure 3.**
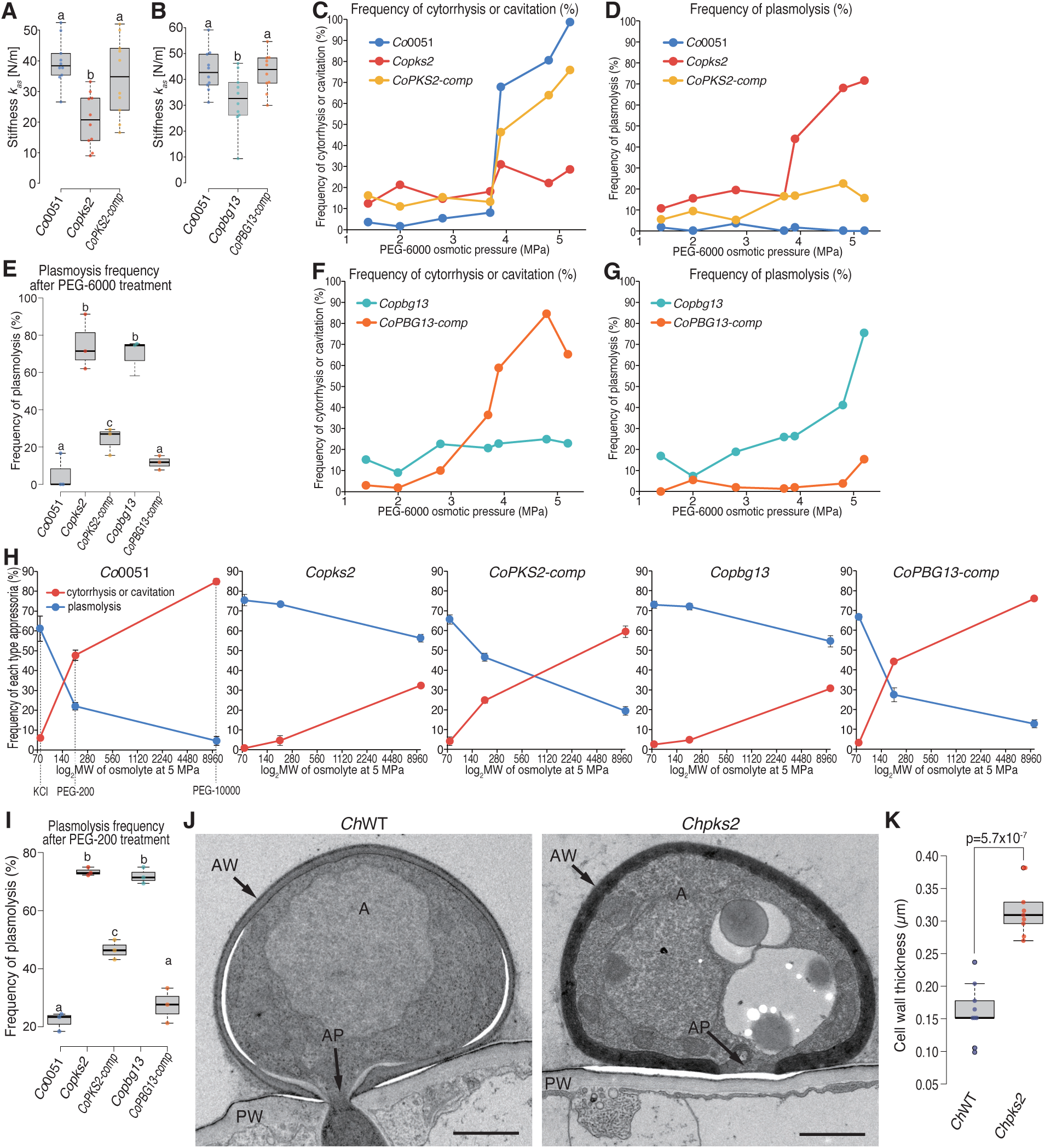
Appressoria of *Copks2* and *Copbg13* exhibit loss of turgor pressure. (**A, B**) Apparent stiffness (*k*_as_) of appressoria was measured using AFM (n=10). Different characters indicate a statistically significant difference (p<0.05, Tukey HSD) (A, B, E, I). (**C, F**) Frequency of cytorrhysis or cavitation in appressoria upon treatment with PEG-6000. Each dot represents frequency from at least 50 appressoria (C-H). (**D, G**) Frequency of plasmolysis in appressoria upon treatment with PEG-6000. Horizontal axes indicate the osmotic pressures of PEG-6000 solutions. (**E, I**) Frequency of plasmolysis in appressoria upon treatment with PEG-6000 (E) or PEG-200 (I) at 5 MPa. Three data points were acquired for each strain. (**H**) Frequency of appressorium response to different osmolytes. Horizontal axes indicate the molecular weight of three osmolytes (KCl, PEG-200, PEG-10000). Each dot represents the mean frequency of three replicates. Error bars: ±SE. (**J**) TEM analysis of median sections through appressoria. A: appressorium, AP: appressorial pore, AW: appressorial cell wall, PW: plant cell wall. Scale bars: 1 µm. (**K**) Measurement of the thickness of appressorial cell walls in TEM images (n=8). p-value was calculated using a two-tailed t test. In A-B and C-I, three and two independent experiments yielded similar results, respectively.

To estimate the pore size of the appressorial cell wall, we used the solute exclusion technique (*37*, *38*), by comparing the molecular weight of osmolytes that could permeate appressorial cell walls. In *Co*0051, treatment with a small osmolyte, KCl (MW: 74 Da), resulted in a higher rate of plasmolysis than cytorrhysis or cavitation (Fig. 3H). Conversely, plasmolysis occurred less frequently with larger molecules, PEG-200 (200 Da) and PEG-10000 (10 kDa), suggesting that *Co*0051 inhibit the entry of molecules larger than PEG-200. In contrast, *Copks2* showed a high frequency of plasmolysis with all three osmolytes, while *CoPKS2*-*comp* showed a lower frequency of plasmolysis with larger osmolytes (Fig. 3H-I), indicating that *Copks2* cannot inhibit the entry of these molecules. Considering the estimated molecular radius of each osmolyte in liquid (K^+^: 0.33 nm, Cl^-^: 0.36 nm, PEG-200: 1.7 nm, and PEG-10000: 5 nm) (*39–41*), pore sizes of *Co*0051 and *Copks2* are expected to be 0.36 to 1.7 nm and >5 nm, respectively. Similar results were obtained with *Copbg13* and *CoPBG13-comp* (Fig. 3H-I). Consistently, transmission electron microscope (TEM) analysis revealed that *PKS2* affects the morphology of the appressorial cell wall (Fig. 3J-K). The *Ch*WT cell wall was thin with a smooth inner surface, while that of *Chpks2* was thick with an uneven inner surface and had a more fibrous structure. In summary, these findings underscore the critical roles of *PKS2* and *PBG13* in maintaining the semipermeable barrier in appressorial cell walls by reducing their pore size.

### PKS2 and PBG13 synthesize DHHA polymers

To identify the SMs produced by PKS2 and PBG13, we generated *Mo* strains over-expressing *PKS2* and/or *PBG13* derived from *Co*, *Ch* and *Mo* (fig. S8A-D). Metabolite analysis of these strains revealed distinct compound peaks in the *OE:MoPKS2A-ChPBG13* strain that were absent in the *Mopks2a* parental strain (fig. S8C). High-resolution mass spectrometry (HRMS) analysis showed that the chemical formula corresponding to the observed series of regularly spaced peaks is (C_6_H_10_O_3_)n. For instance, P-1041, P-1171, and P-1301 correspond to (C_6_H_10_O_3_)_8_, (C_6_H_10_O_3_)_9_, and (C_6_H_10_O_3_)_10_, respectively, with each differing by one C_6_H_10_O_3_ monomer unit, exact mass 130.06 (Fig. 4A). These results demonstrate that *Mo*PKS2A and *Ch*PBG13 synthesize polymers of C_6_H_10_O_3_. As typical fungal PKSs employ acetyl-CoA as a starter unit and malonyl-CoA as an extension unit (*42*), the predicted monomer structure, represented as predicted PKS2 product (**1**) (fig. S8E), likely forms ester-linked polymers (C_6_H_10_O_3_)n. To verify this, we treated the polymers with alkali to cleave the ester bond, hypothesizing that the resulting monomers would correspond to compound **2** (3,5-dihydroxyhexanoic acid or DHHA), which would then cyclize into cyclized monomer (**3**) (4-hydroxy-6-methyltetrahydropyran-2-one) upon acidification (Fig. 4B). Indeed, HRMS and 1D and 2D nuclear magnetic resonance (NMR) analysis confirmed that the predicted structure of **3** matched the structure of the compound purified after alkali and acid treatments of *OE:MoPKS2A-ChPBG13* cultures (Fig. 4C, fig. S9). In addition, alkaline-treated extracts from *Ch*WT appressoria yielded a specific peak with an exact mass of 130.06, corresponding to cyclized monomer (**3**), which was not detected in the alkaline-treated extracts of *Chpks2* and *Chpbg13* appressoria (Fig. 4D). These findings document that PKS2 and PBG13 biosynthesize DHHA polymers, corroborating the enzymatic functionality of these proteins in SM production.

**Figure 4.**
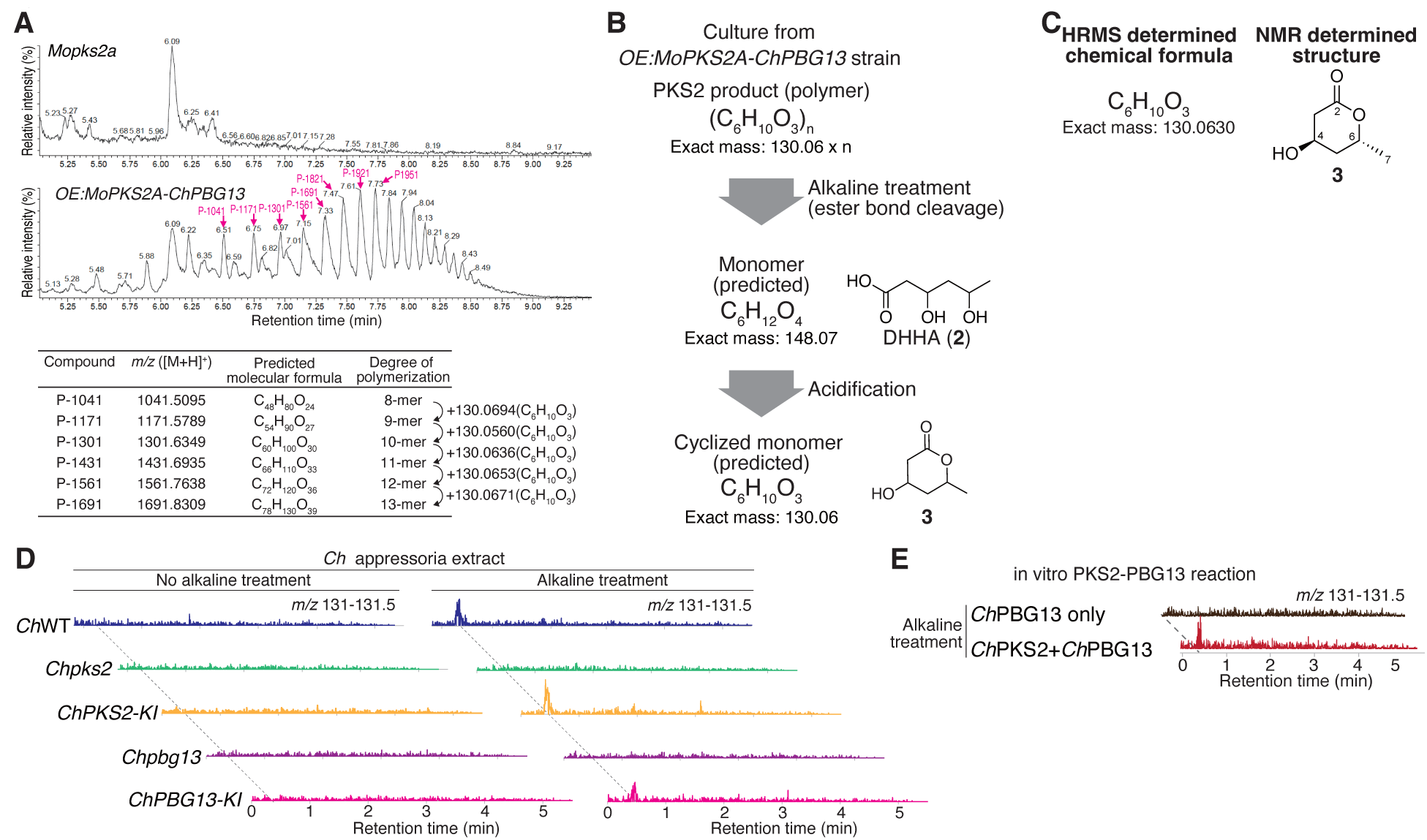
PKS2 and PBG13 produce polymers with a monomer unit of exact mass 130.06. (**A**) HRMS analysis of the *OE:MoPKS2A-ChPBG13* strain identified compound peaks corresponding to polymers with monomer units having an exact mass of 130.06. (**B**) Predicted structure or chemical formula of the PKS2-product in polymer form, indicating the monomer unit after alkaline-treatment (**2**: 3,5-dihydroxyhexanoic acid, or DHHA) and the cyclized monomer upon acidification (**3**: 4-hydroxy-6-methyltetrahydropyran-2-one). (**C**) HMRS and NMR analysis determined that the chemical formula and structure of PKS2-monomer matched **3** exactly. Details on metabolite preparation and structure elucidation are described in fig. S9. (**D**) A metabolite with a molecular weight of 130, corresponding to cyclized monomer (**3**), was detected in the appressoria extract. (**E**) In vitro reaction of *Ch*PKS2 and *Ch*PBG13 proteins with substrates, acetyl-CoA and malonyl-CoA, produced a metabolite with a molecular weight of 130, corresponding to cyclized monomer (**3**).

Finally, we tested whether *Ch*PKS2 and *Ch*PBG13 are sufficient to synthesize DHHA through in vitro biochemical reactions. First, we engineered strains by knocking-in the Twin-Strep-tag (TStag) coding sequences upstream of the stop codons of the native *ChPKS2* and *ChPBG13* (*ChPKS2-TStag-KI* and *ChPBG13-TStag*-*KI*). These strains retained pathogenicity, demonstrating that the tagged proteins are functional, and we then affinity-purified those proteins from appressoria (fig. S10A-C). The reaction mixture of the purified tagged proteins with acetyl-CoA and malonyl-CoA as substrates was subjected to alkaline treatment (Fig. 4E) and a peak with a molecular weight of 130 was identified, confirming that *Ch*PKS2 and *Ch*PBG13 are necessary and sufficient for DHHA production.

## Discussion

Until now, melanin was recognized as the sole biochemical component forming the semipermeable barrier in fungal appressoria. Our study identified two novel enzymes, PKS2 and PBG13, essential for the semipermeability of appressorial cell walls, and demonstrated their role in the biosynthesis of a DHHA polymer (fig. S11A). Previous reports on melanin (*18*, *43*) and our findings suggest that melanin and the PKS2-PBG13 products confer different properties to the appressorial cell wall. While DHN-derived melanin provides physical rigidity to withstand high turgor pressures (*18*, *43*), the DHHA-derived polymers confer semipermeability, preventing osmolyte permeation. The mechanism by which DHHA products enhance semipermeability remains to be fully elucidated; however, we propose two potential models: either DHHA reduces the pore size of the cell wall by interacting with components such as polysaccharides (chitin, glucans) and melanin, or it polymerizes independently to fill the spaces between these components, thereby reducing pore size (fig. S11B).

Future exploration of the nanoscale physical structure of appressorial cell walls and the interactions of DHHA polymers with other cell wall components will provide critical insights. Given that chemical inhibitors of melanin biosynthesis have been pivotal in controlling appressorial fungal pathogens (*44*, *45*), our findings suggest that PKS2 and PBG13 may provide additional molecular targets for crop protection. This is particularly urgent as the threat to global food security from pathogens, such as wheat blast, is expanding due to climate change (*46*, *47*).

## Acknowledgments

We appreciate Drs Takayoshi Awakawa and Uta Paszkowski for critical feedback on the manuscript. We thank Akiko Ueno, Takuma Aita, Hayato Fukuda, Reika Hiraishi, and Fusuke Matano for their technical assistance. English editing was supported by ChatGPT-3.5.

## Author contributions

Conceptualization: N.K., K.Sh.; methodology: N.K., T.M., K.Mi. T.N., R.O.; formal analysis: N.K., T.M., K.Mi, T.N., H.K., R.O.; investigation: N.K., T.M., K.Mi, T.N., K.Y., K.Sa., N.I., K.Ma., P.G., H.K., R.O.; resources: H.K., T.F., R.O., K.Sh.; writing-original draft: N.K., T.M., K.Mi., T.N., P.G., R.O. ; writing-review and editing: N.K., R.O., K.Sh.; visualization: N.K., M.T., K.Mi., T.N.; supervision: H.K., T.F., R.O., K.Sh.; funding acquisition: N.K., M.T., K.Sh.

## Data and materials availability

All data supporting the findings of this study are included within the article or supporting materials. Original raw data for all experiments will be provided upon request. The genome sequence used in this study is available from the NCBI BioProject under the accession number PRJNA171217. The RNA-seq data can be accessed from the NCBI Gene Expression Omnibus with the ID GSE178879. The wild-type organisms used in this study are available from the respective resource centers or companies that provided them. The methods for creating the genetically modified derivative organisms based on these wild-type organisms are described in this paper. Importation to the requester’s country requires compliance with respective quarantine procedures.

## Supplementary Materials

### Materials and Methods

#### Fungal strains, culture, and storage

*Colletotrichum orbiculare* 104-T (*48*) (NARO Genebank ID: MAFF 240422), *Colletotrichum higginsianum* IMI349063A (*29*, *49*), *Magnaporthe oryzae* Kita1 (NARO Genebank ID: MAFF 101512) and 70-15 (*50*) strains were used as wild type (WT) in this study. *Co*0051, a *Coura3a* mutant strain, demonstrating pathogenicity similar to the *Co*WT strain (*27*), was also employed as a positive control for pathogenicity. *Co*, and *Ch* strains were grown on PDA (Nissui) and Mathur’s media (*51*) agar (MA) (16 mM glucose, 5 mM MgSO_4_, 20 mM KH_2_PO_4_, OXOID mycological peptone 0.22% [w/v], agar 3% [w/v]), respectively. *Mo* strains were cultured on PDA (Difco Co.), OMA (Difco Co.), YG medium (0.5% yeast extract, 2% glucose), or OM medium (6% oatmeal [Quaker]) at 25 °C or 28 °C. All strains were stored in 25 % (w/v) glycerol at −80 °C for long-term preservation.

#### Prediction of *Co*SMKGs and BGCs

BGCs and SMKGs were predicted from the *Co* 104-T genome sequence (NCBI accession: PRJNA171217) (*23*) using three software tools: antiSMASH 4.0 (*52*), SMIPS (*53*), and SMURF (*54*). Candidate BGCs were tentatively defined as contiguous regions of the *Co* genome meeting the following criteria: containing at least two genes predicted to be part of a cluster by one or more of the software tools used and harbouring at least one SMKG listed in table S1. Details regarding all predicted SMKGs and BGCs are in tables S1 and Data S1, respectively.

#### RNA-seq analysis

Previously reported RNA-seq data for the *Co* 104-T strain were utilized, sourced from the Gene Expression Omnibus of NCBI (accession number: GSE178879) (*24*). Briefly, RNA-seq samples included spore (SP), vegetative hyphae (VH), and *Nicotiana benthamiana* leaves infected at 1 DPI, 3 DPI, and 7 DPI. All samples were analyzed in three biological replicates. The average log counts per million mapped reads (CPM) and average reads per kilobase of transcript per million mapped reads (RPKM) values for *Co* genes are summarized in Data S2. Raw reads were aligned to the *Co* whole genome using STAR (*55*) (v2.6.0a) with the parameter ’--alignIntronMax 1000’, followed by quantification of transcript counts using Rsubread (*56*). RPKM values were calculated using the rpkm function in edgeR (*57*) after normalization with the TMM normalization procedure. The average RPKM values for each sample type at each lifecycle stage were summarized across three biological replicates using the rpkmByGroup function. CPM counts were calculated using the edgeR cpm function on the TMM normalized edgeR DGEList object, filtered by expression. To calculate average logCPM values, biological replicates of each sample type were subsetted, and the aveLogCPM function was applied to each subset. To assess the relative expression levels of each gene across samples, we employed the following method: the CPM values of SP, VH, 1 DPI, 3 DPI, and 7 DPI were added together, and then each sample’s CPM was normalized by this total to yield a relative expression value, with the highest expression set to 1. The identification of *Co*SMKGs upregulated at 1 DPI, 3 DPI, and 7 DPI was conducted through the following methodology: Initially, all 73 *Co*SMKGs were clustered based on their relative expression levels in each sample using the pheatmap package on R (https://www.R-project.org/). Subsequently, a subset of SMKGs exhibiting specific up-regulation at 1 DPI, 3 DPI, and 7 DPI was isolated. Within this subset, SMKGs upregulated at 1 DPI were further categorized based on their absolute expression values, quantified as RPKM. Similarly, groups of SMKGs displaying peaks in expression at 3 DPI and 7 DPI underwent analogous sorting procedures. Heatmaps representing relative and absolute expression values for each gene were drawn using the ‘pheatmap’ function in R.

#### Search for homologs of PKS2 and PBG13

PKS2 and PBG13 homologs were surveyed from 19 fungi, including eight species from the *Colletotrichum* genus, 10 species of other plant pathogenic fungi (*2*), and a saprophyte (*Neurospora crassa*). *Co*PKS2 (Genbank ID: TDZ25780.1) and *Co*PBG13 (Genbank ID: TDZ25781.1) were used as queries, and the NCBI BlastP searches were performed against the 19 species with default settings. Proteins with a query cover of 90% or more and a partial identity of 50% or more were considered homologs. Details of each homolog are listed in table S2.

#### DNA construction

*Co*, *Ch*, and *Mo* genomic DNAs used as templates for PCR were isolated by DNeasy Plant Mini Kit (QIAGEN). PCR was performed using either KOD One PCR Master Mix -Blue-(Toyobo), KOD Plus Neo (Toyobo), or KOD FX Neo (Toyobo) DNA polymerases. Plasmids and DNA fragments for fungal transformation were constructed using In-Fusion HD Cloning Kit (Takara) or NEBuilder HiFi DNA Assembly Master Mix (NEB). The constructions for the *CoURA3*-based marker recycling method were designed as described in our previous report (*27*). DNA constructs and primers used are listed in Data S3 and S4, respectively.

#### Fungal transformation

Transformation of *Co* and *Ch* strains with or without the CRISPR-Cas9 system followed established procedures as described in previous reports (*25*, *30*)*. Mo* was transformed using *Agrobacterium tumefaciens*-mediated transformation (ATMT) (*58*), employing *A. tumefaciens* strain C58 (*59*, *60*). *Mo* transformants were selected using either 500 µg/ml hygromycin B (Wako Pure Chemical) or 200 µg/ml blasticidin S (Wako Pure Chemical). Details of *Colletotrichum* (including *Co* and *Ch*) and *Mo* strains were provided in Data S5 and S6, respectively. Information regarding the gRNAs used is listed in table S3.

#### Spore preparation

*Co* strains were grown on 20 ml PDA in a 90 mm plastic Petri dish or 100 ml PDA in a 300 ml flask for six days at 25°C in darkness to induce sporulation (*25*). The resulting spores were then harvested, filtered through a cell strainer (100 µm pore-size, Corning), and suspended in sterilized water. *Ch* strains were cultured on MA in a 250 ml Erlenmeyer flask, covered with aluminium foil and sponge plugs for aeration, and incubated at 25 °C in darkness to induce sporulation (*30*). After 7 to 9 days, the spores were harvested and suspended in sterilized water. *Mo* strains were cultured on OMA plates at 28°C for 7 days. Aerial hyphae were removed, and spore formation was induced under BLB light (FL20S BLB, Toshiba Co.) for 4 days. The formed spores were suspended in 0.02% (w/v) Tween 20 using a drawing brush, filtrated through a gauze, and collected by centrifugation (1000xg, 10 min). The spores were then suspended in 0.02% (w/v) Tween 20. The concentration of spores was determined using a hemacytometer.

#### Fungal inoculation to monitor disease symptoms

Inoculation of *Co* onto cucumber was conducted following a previously established protocol (*27*). Briefly, spores were suspended in sterilized water at a concentration of 5×10^5^ spores/ml. Seeds of *Cucumis sativus* L. Suyo strain (cucumber) (Sakata Seed Corp.) were sown on a soil mixture comprising an equal amount of SupermixA (Sakata Seed Corp.) and vermiculite (VS Kakou). The cucumbers were then incubated in a growth chamber at 24°C under a 10h-light/14h-dark photoperiod. Cotyledons were harvested with scissors at 6-9 days post germination (DPG), and 5 µl droplets of the spore suspension were inoculated onto each cotyledon using a pipette. Inoculated leaves were placed in a humid chamber and incubated under the same conditions for cucumber growth. At 5-7 DPI, photographs of disease spots were captured using a Sony α7III camera equipped with a macro lens (FE 50mm F2.8 Macro) and a copy stand.

Inoculation of *Ch* onto *A. thaliana* Col-0 leaves was performed as follows (*30*). *A. thaliana* Col-0 grown at 22 °C under an 10-h light photoperiod for 35 days was used for inoculation. A 10 µl droplet of *Ch* spore suspension at a concentration of 1×10^6^ spores/ml in sterilized water was inoculated onto true leaves of *A. thaliana* Col-0 and incubated under the same conditions as the plant growth in a humid box. At 7 DPI, photographs were taken as described for *Co* inoculation.

Inoculation of *Mo* onto the susceptible rice cultivar Nipponbare (Nouken Corp.) was performed as follows. A spore suspension at a concentration of 1×10^5^ spores/ml in 0.02% Tween 20 was sprayed onto rice leaves at the three-leaf stage using an airbrush. The inoculated leaves were then incubated in a wet box at 25 °C for 17 h. Subsequently, the sprayed plants were cultured in controlled-environment chambers under a 14-h light photoperiod for 6 d and photographed. For each *Mo* strain, 10-14 plants were used.

#### *Co* invasion frequency on cucumber

Experiments were carried out following a previously described method (*26*). Spore suspension of *Co* was adjusted to a concentration of 1×10^5^ spores/ml in distilled water. Cucumbers were sown and cultivated according to the method described in the “Fungal inoculation to monitor disease spots” section. At 6 DPG, cotyledons were detached with scissors and placed on a moistened Kimwipe in a plastic Petri dish (Corning), with the abaxial side facing upwards. For each cotyledon, two droplets of 5 µl spore suspension were inoculated. The plastic dishes containing the inoculated leaves were covered with lids, sealed with plastic wrap to maintain humidity, further covered with aluminium foil, and then incubated at 24°C. At 65 HPI, the inoculated leaves were stained with trypan blue following previously described methods with modifications (*61*, *62*) to visualise *Co* invasive hyphae in cucumber leaves. In brief, inoculated leaves were boiled in lactophenol ethanol trypan blue solution (25% [v/v] lactic acid, 25% [w/v] phenol, 50% [v/v] ethanol, 0.2 mg/ml trypan blue, 25% [v/v] glycerol) for 5 min in a water bath and then destained in a 2.5 g/ml chloral hydrate solution overnight. Trypan blue-stained invasive hyphae emerging beneath the appressoria were counted using a Nikon Eclipse Si microscope. The invasion frequency was calculated as the percentage of appressoria that produced invasive hyphae out of the total number of observed appressoria.

#### Morphology of spore, colony, and appressorium

*Co* and *Ch* samples (spore and colony) were prepared as described in previous reports (*26*, *30*). Appressoria and spores of *Co* and *Ch* were observed and photographed using microscopy (Olympus) and a LP74 microscope digital camera (Olympus). The UPlanSApo 100x objective lens (Olympus) was used for differential interference contrast (DIC).

#### Quantitative PCR

RNAs were extracted from spores of *Ch*WT and in vitro induced appressoria at various time points ranging from 5 to 48 HPI. The in vitro appressoria were induced following the procedure outlined in “Subcellular localization of *Ch*PKS2 and *Ch*PBG13”. At each time point, in vitro appressoria were scraped with 0.6 ml of chilled 1-Thioglycerol/Homogenization solution from the Maxwell RSC Plant RNA Kit (Promega) and the mixture was stored at −80 °C until RNA isolation. To obtain RNAs from spores serving as the “0 HPI” control, the spore suspension was first filtered through an MF-Millipore membrane (0.22 µm pore size, Millipore). The filtered spores were scraped from the membrane, frozen using liquid nitrogen in a 2-ml steel top tube with a 5 mm diameter zirconia bead, crushed using a shaker (ShakeMaster NEO [BMS]), and stored at −80°C until RNA isolation. RNAs were isolated from both in vitro appressoria and spore samples with genomic DNA removal using the Maxwell RSC Plant RNA Kit (Promega) and the Maxwell RSC 48 instrument (Promega), following the manufacturer’s instructions. Reverse transcriptions were conducted using the RevTra Ace qPCR RT Kit (Toyobo) with a Primer Mix containing random primers and oligo dT following the manufacturer’s instructions. Quantitative PCR (qPCR) was carried out using THUNDERBIRD Next SYBR qPCR Mix (Toyobo) and an MX3000P Real-Time qPCR System (Stratagene). Primers used are listed in Data S4.

#### Subcellular localization of *Ch*PKS2 and *Ch*PBG13

Spores of *Ch* strains were harvested and suspended in distilled water at 2×10^6^ spores/ml. The spore suspension was applied onto a glass-bottom dish (ibidi) using a disposable hand spray. The dishes were then placed in a humid box at 25°C in darkness. At 8 HPI, appressoria formed on the dishes were observed using a confocal microscope, Leica TCS SP8 X, equipped with a White Light Laser. For immunoblotting of *Ch*PKS2-mGreenLantern and *Ch*PBG13-mScarlet3, in vitro appressoria were induced following a previously described method (*30*). Briefly, a 40 ml spore suspension of *Ch* strains at a concentration of 2×10^6^ spores/ml was poured into a plastic dish (14×10 cm). After a 40 min incubation to allow spores to settle on the bottom surface, a piece of sterilized nylon mesh (50 µm pore size, 13x9 cm) covered the water surface, and the water was carefully discarded, leaving the nylon mesh on the bottom of the plate to keep spores moist. The plate was then incubated at 25°C in darkness within a humid box to induce appressoria formation. At 8 HPI, 0.5 ml protein extraction buffer (150 mM NaCl, 1% [w/v] Igepal CA-630, 5% [w/v] glycerol, 50 mM Tris-HCl pH8.0, 2 mM TCEP) with Protease Inhibitor Cocktail for plant cell and tissue extracts (Sigma-Aldrich) at 100 times dilution was added to the plate after removing the nylon mesh, and appressoria were harvested using a cell scraper. The cell lysate was centrifuged (5 min, 10,000 g, 4 °C) and the supernatant was mixed with LDS sample buffer (Thermo Fisher Scientific) for subsequent SDS-PAGE and immunoblotting analysis. Anti-GFP antibody (ab290, abcam) and anti-mCherry antibody (ab167453, abcam), and anti-rabbit antibody-peroxidase conjugated (NA934V, Cytiva) were used. Chemiluminescence was induced using SuperSignal West Femto Maximum Sensitivity Substrate (Thermo Fisher Scientific), and the luminescence was detected by a LAS4000 imager system (GE Healthcare).

#### AFM measurements of in vitro appressoria

Spore suspensions of each *Co* strain, adjusted at 2×10^6^ spores/ml in distilled water, were applied onto polystyrene bottom dishes (TPP) with a 40 mm diameter using a disposable hand spray. These dishes were then placed in a humid box at 25°C in darkness. At 20-22 HPI, appressoria formed on the dishes were immersed in ultra pure water. Atomic force microscopy (AFM) measurement was performed using a JPK Nanowizard4 (Bruker Nano GmbH) Quantitative Imaging (QI) mode at room temperature (25°C). For the height image measurement in fig. S6D, a silicon cantilever with a nominal spring constant of 26 N/m (160AC-NG, OPUS, MikroMasch) was used. The force setpoint value was 100 nN, and the measured size was 15.4×15.4 μm^2^ with 256×256 pixels. The Z length and vertical tip scanning speed were 2.5 μm and 250 μm/s, respectively. For measurement of apparent stiffness (*k*_as_) in Fig. 3A and B the detailed procedure is described in fig. S6. The tip apex was laterally cut using a focused ion beam (HeliosG4 CX, Thermo Fisher Scientific). A silica bead (diameter=1.86 μm, SiO2MS-2.0, Cospheric) and glue (353ND, Epoxy Technology) were then attached using a micromanipulator (AxisPro, Microsupport), with the glue being fixed by heating at 150°C for 30 min (fig. S6A-B). To measure the stiffness of in vitro appressoria, the whole structure of the appressorium and spore was first measured by AFM. Subsequently, a region of interest (ROI) above the appressorium with a size of 3.5×3.5 μm^2^ was selected. In QI mode measurement within the selected ROI area (fig. S6C-D), the force setpoint value was 1,000 nN, and the measured size was 3.5×3.5 μm^2^ with 16×16 pixels. The Z length and vertical tip scanning speed were 1,000 μm and 8.33 μm/s, respectively. A single force curve was obtained above each appressorium, and the appressorium stiffness (*k*_as_) was calculated from the slope of the force curve fitted with a linear function in the region where the interaction force increases linearly with the indentation length, based on elastic shell theory (*35*) (fig. S6E). Ten appressoria were analyzed for each strain (n=10, fig. S6F).

#### Incipient cytorrhysis assay

Appressorial turgor was indirectly estimated following a previously established method(*15*). Spore suspension was adjusted to a concentration of 2×10^6^ spores/ml in sterilized water. Spores were placed in tiny water droplets using a disposable hand spray on 1 x 1 cm pieces of a hydrophobic membrane (PTCEF, aclar film), which allows appressoria development. The inoculated membranes were then placed in a humid box and incubated in darkness at 25 °C. At 20-21 HPI, tiny water droplets were replaced with PEG-6000 (Sigma-Aldrich) solutions at various concentrations: adjusted to 20, 30, 35, 37.5, 40, 45, and 50% (w/v) in sterilized water. The osmotic pressure of each PEG-6000 solution was estimated to be 1.4, 2, 2.8, 3.7, 3.9, 4.8, and 5.2 MPa, based on a previous report (*63*). After incubating 10 min in PEG solution, the frequency of appressoria cytorrhysis or cavitation, which is evidence of the solute exclusion (*36*) (fig. S7), was measured from at least 50 cells in each concentration of PEG-6000 using phase-contrast microscopy (BX51, Olympus). In parallel, the frequency of plasmolysis, which occurs only when the solute molecules, PEG-6000, diffuse through the cell wall (fig. S7), was determined.

#### Solute exclusion technique

Appressorial cell wall pore size of *Co* strains was estimated using a solute exclusion technique (*15*, *37*, *38*). Appressoria were prepared using the method described in the section “Incipient cytorrhysis assay”, and treated with an aqueous solution of osmolytes of various molecular sizes including KCl (74.6 Da), PEG-200 (200 Da), and PEG-10000 (10 kDa), for 10 min. The osmotic pressure of the solutions was adjusted to 5 MPa based on a previous report (*63*). The frequencies of plasmolysis, cytorrhysis, and cavitation were measured from at least 50 appressoria. Plasmolysis occurs only when a solute can pass through the pores in the cell wall, while cytorrhysis or cavitation occurs when the solute is excluded by the cell wall (fig. S7). Therefore, the threshold for molecule size that can pass through the cell wall lies at the smallest molecular size where the frequency of cytorrhysis or cavitation is higher than the frequency of plasmolysis.

#### TEM analysis

*A. thaliana* Col-0 seeds were sown in a peat-based compost (Floradur-B, Floragard, Oldenburg, Germany), and grown in a Percival AR-36L3 growth chamber (CLF PlantClimatics GmbH, Wertingen, Germany; 12-h photoperiod, 230 μmol m^-2^ s^-1^ photon flux density, 23/21°C day/night temperature, 70% relative humidity). The cotyledons of 12-day-old seedlings were inoculated with 2 µl droplets of *Ch* spore suspension (5 x 10^5^ spores/ml), placed inside plastic propagators to maintain 100 % humidity, and incubated in a growth chamber (25 °C, 12h photoperiod cycle, 80 μmol m^-2^ s^-1^ photon flux density). At 40 HPI, the cotyledons were harvested using a razor blade and vacuum-infiltrated for 10 min with a fixative solution containing 2.5% (v/v) glutaraldehyde in sodium cacodylate buffer (0.05 M, pH7.2). After fixation for 3 h at room temperature with continuous agitation on a rotator, the samples were rinsed in 0.05 M cacodylate buffer (three changes, 10 min each), and fixed in 1% (w/v) osmium tetroxide in 0.05 M cacodylate buffer for 2 h. After rinsing in deionized water (three changes, 10 min each), the samples were dehydrated through an ethanol series: 25% (30 min), 50% (30 min), 70% (1 h), 90% (1 h), and 100% ethanol (two changes, 1 h each). After transfer to a 1:1 mixture of ethanol and acetone for 30 min, then 100% acetone (two changes,15 min each), the samples were infiltrated with increasing concentrations of Agar LV epoxy resin (hard grade, Agar Scientific, Stansted, UK) in acetone: 20% (30 min), 40% (30 min), 60% (1h), and 80% (1h). The samples were then infiltrated with 100% resin (two changes per day for 3 days), transferred to flat embedding moulds and polymerized at 65 °C for 18 h. Appressoria were identified by light microscopy and prepared for targeted sectioning as described previously (*64*). Ultrathin sections (90 nm) were mounted on Formvar-coated copper slot grids and stained with Uranyless^©^ (1 min, Delta Microscopies, Mauressac, France) and lead citrate (1 min), before observation with a JEOL JEM-1400 TEM operating at 80 kV.

#### Metabolite analysis

Fungal cultures, including mycelia and culture broth, were treated with four volumes of ethanol, and the supernatant was dried under nitrogen. The dried material was dissolved in 80% ethanol and subjected to analysis. Fungal metabolites were analyzed via ultra-performance liquid chromatography (UPLC)/mass spectrometry (MS) using the Waters Acquity UPLC H-Class-QDa system (Waters, Milford, MA, USA). A reversed-phase column (BEH C18, 2.1 × 50 mm, 1.7 µm, Waters) was used at a flow rate of 0.6 mL/min. The gradient elution was achieved by varying the composition of acetonitrile (solvent B) and 0.1% formic acid-water (solvent A) as follows: starting with 5% B from 0 to 1 min, increasing to 95% B from 1 to 4.5 min, holding at 95% B from 4.5 to 5.5 min, and re-equilibrating with 5% B from 5.5 to 8 min. Detection of metabolites was carried out using positive ion electrospray ionization (ESI).

#### Alkaline treatment of metabolites

The fungal culture, including mycelia and culture broth, underwent alkaline treatment by adding 0.1 volume of 10 N potassium hydroxide, followed by a 2-hour incubation at 25 °C. The solution was acidified by adding 0.1 volume of 12 N hydrochloric acid, treated with 4 volumes of ethanol, and the resulting supernatant was dried under nitrogen. The dried material was dissolved in 80% ethanol, neutralized with 1 N potassium hydroxide, and analyzed.

#### Isolation and structural identification of metabolite by NMR

The metabolite isolation process is summarized in fig. S9A. Briefly, 50 ml of the hydrolyzed culture, as described in the section “Alkaline treatment of metabolites”, was adjusted to a pH of 1 to 2 using 1 N HCl. The solution was then extracted with ethyl acetate (EtOAc) to remove lipophilic materials. The aqueous phase was further extracted with 1-butanol (1-BuOH), and the 1-BuOH solution was evaporated to yield 98.6 mg of a polar extract. This extract was subjected to column chromatography on Sephadex LH-20 and eluted with 100% MeOH to yield four fractions. The third fraction (F003) was separated by HPLC on Capcell Pak ADME (10 mm id x 250 mm, 5 µm, OSAKA SODA, Osaka, Japan), resulting in 3.8 mg of a crude compound **3,** a colorless amorphous solid (F007), which was used for structural determination.

The molecular formula of compound **3** was determined to be C_6_H_10_O_3_ by high-resolution electrospray ionization time-of-flight mass spectrometry (HR-ESI-TOFMS), with an observed *m/z* value of 131.0710 [M+H]^+^ and a calculated value for C_6_H_11_O_3_ 131.0708 (Synapt G2, Waters, MA, USA). The structure of compound **3** was elucidated using 1D and 2D nuclear magnetic resonance (NMR) spectroscopy, including HMQC, edited-HMQC, DQF-COSY, HMQC-TOCSY, and HMBC spectra (JMN-ECZ600, JEOL, Tokyo, Japan) (fig. S9B). The relative stereostructure was determined by nuclear Overhauser effect spectrometry (NOESY) correlation (fig. S9C-D).

#### In vitro reaction of *Ch*PKS2-TStag and *Ch*PBG13-TStag proteins

Lysates from in vitro induced appressoria of *ChPKS2-TStag-KI* and *ChPBG13-TStag-KI* strains were prepared according to the method described in the section “Subcellular localization of *Ch*PKS2 and *Ch*PBG13“, with modifications for scale-up. Briefly, appressoria were induced in 22.5x22.5 cm square plastic dishes (BioAssay Dish, Thermo Fisher Scientific) covered with 22x22 cm sterilized nylon meshes. A 10 ml protein extraction buffer containing a protease inhibitor cocktail was used for each dish. Insoluble debris in appressorial lysates was removed by centrifugation four times (4 °C, 10,000 g, 3 min) and filtration through a 0.22 µm PES membrane (Sartolab 180C4, Sartorius). The resulting supernatant was loaded onto a chromatography column (StrepTrap XT, Cytiva) pre-equilibrated with protein extraction buffer. The column was washed subsequently with protein extraction buffer without a protease inhibitor cocktail and wash buffer (50 mM NaCl, 5% [w/v] glycerol, 50 mM Tris-HCl pH8.0, 2 mM TCEP), and the bound protein were eluted with wash buffer supplemented with 50 mM biotin. The eluted fractions containing *Ch*PKS2-TStag or *Ch*PBG13-TStag were pooled and buffer-exchanged into 50 mM NaCl, 5% (w/v) glycerol, 2 mM TCEP, and 50 mM Tris-HCl pH7.5. The purified *Ch*PKS2-TStag or *Ch*PBG13-TStag proteins were stored at −80 °C until use. A reaction mixture containing 0.5 µM *Ch*PKS2-TStag and 0.15 µM *Ch*PBG13-TStag in 100 µl of 50 mM Tris-HCl (pH7.5), 2 mM MgCl2, 2 mM NADPH, 2 mM TCEP, 2 mM malonyl-CoA (MERK), and 2 mM acetyl-CoA (MERK) was incubated overnight at 25 °C on a rotator at 300 rpm. The reaction mixture without *Ch*PKS2-TStag was used as a negative control. Following incubation, the reaction mixture was treated with alkaline as described in “Alkaline treatment of metabolites” and analyzed by UPLC/MS.

For the detection of *Ch*PKS2-TStag and *Ch*PBG13-TStag by immunoblotting, anti-Strep-tag II antibody (ab76949, Abcam) and anti-rabbit antibody-peroxidase conjugate (NA934V, Cytiva) were used, following the method described in the “Subcellular localization of *Ch*PKS2 and *Ch*PBG13” section.

#### Statistics and data presentation

All statistical analyses were performed using R version 4.3.0 or earlier (https://www.r-project.org/). All graphs were visualized using either R or Excel (Microsoft). Each element in a box plot represents the following: the box shows the interquartile range (IQR), with a horizontal line inside indicating the median. Whiskers extend from the box to the minimum and maximum values within 1.5 times the IQR.

#### Accession numbers

The accession numbers of genes used in this study are listed in table S4.

---

**Fig. S1.**
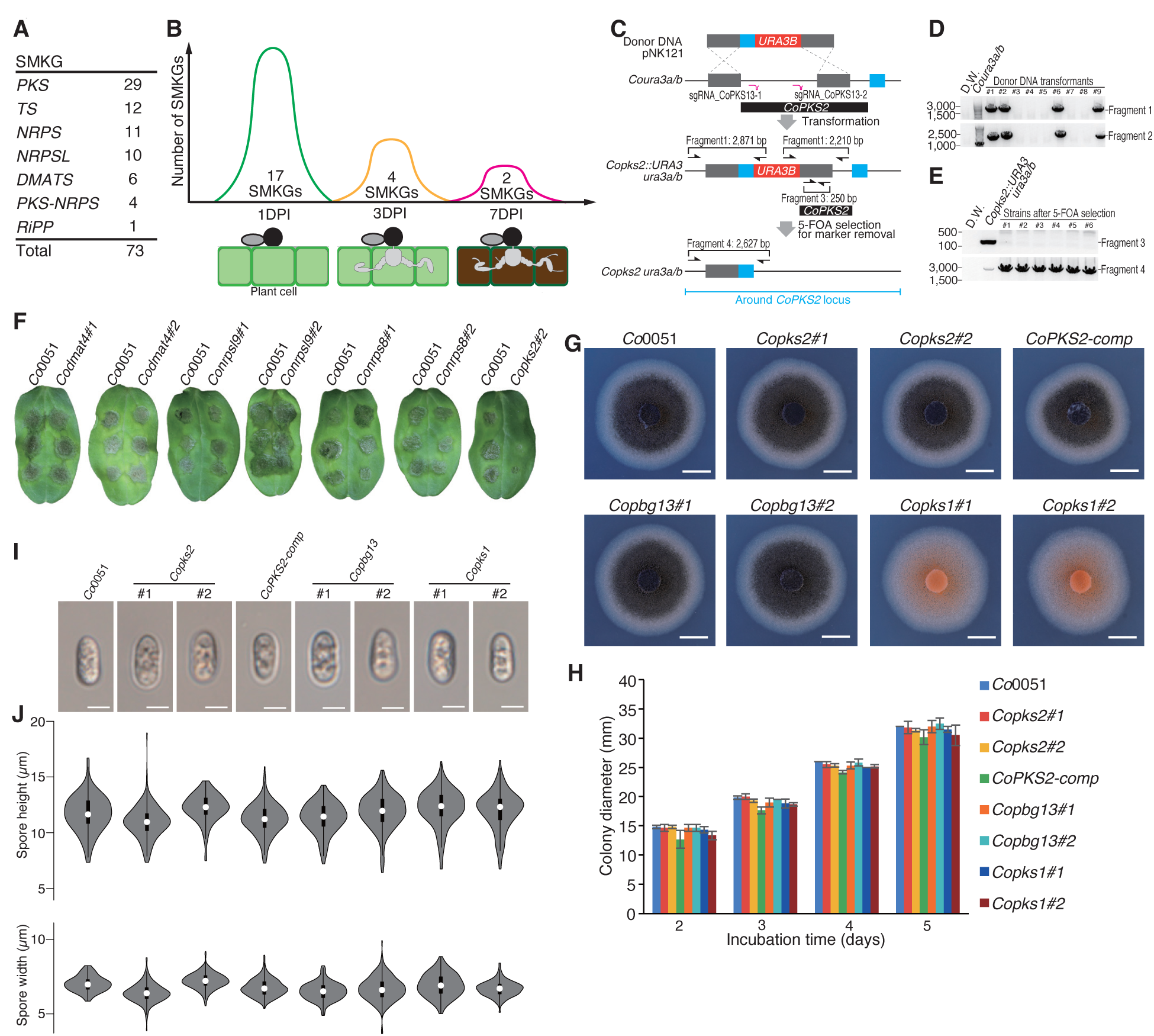
*CoPKS2* and *CoPBG13* knockouts do not affect colony and spore morphology. **(A)** Number of predicted SMKGs in *Co*. **(B)** Number of *Co*SMKGs up-regulated at 1 DPI, 3 DPI, and 7 DPI. The schematics illustrate representative cell status at each infection stage. At 1 DPI, melanized appressorium is formed from spore (oval structure on left). At 3 DPI, invasive hyphae emerge from the base of *Co* appressoria to penetrate living plant cells, marking the biotrophic phase. At 7 DPI, host plant cells surrounding fungal cells are dead (dark brown), indicating the necrotrophic phase. **(C)** Schematics of *CoPKS2* knockout. The donor DNA has a selection marker, *URA3B*(*24*) (red box), flanked by two homology arms (grey boxes). Green boxes represent homology arms to remove *URA3B* from the genome of *Copks2::URA3 ura3a/b* via homologous recombination after 5-FOA treatment. The black box indicates the coding sequence of *CoPKS2*. *Coura3a/b*(*24*), which shows uridine auxotrophy, was used as a parental strain. Magenta lines represent gRNAs of CRISPR-Cas9. (**D, E)** PCR screening of *Copks2::URA3 ura3a/b* and *Copks2/ura3a/b* strains. The primer sets used are depicted in (C). Strains in which bands corresponding to both Fragment1 and Fragment2 were observed were selected as *Copks2::URA3 ura3a/b* strains (D). Strains that exhibited bands corresponding to Fragment 4 but not Fragment 3 were selected as *Copks2/ura3a/b* strains (E). (**F**) Disease symptoms on cucumber leaves caused by *Co* mutant strains. The same experiments as depicted in Fig. 1B was performed. (**G, H**) Colony morphology (G) and diameter (H) of each *Co* strain grown on PDA. Error bars indicate ±SE (n=3). Scale bars represent 5 mm. (**I, J**) Spore morphology (I), height, and width (J) of each *Co* strain. Error bars represent 5 µm. Violin plots were based on measurements of at least 200 spores for each strain.

**Fig. S2.**
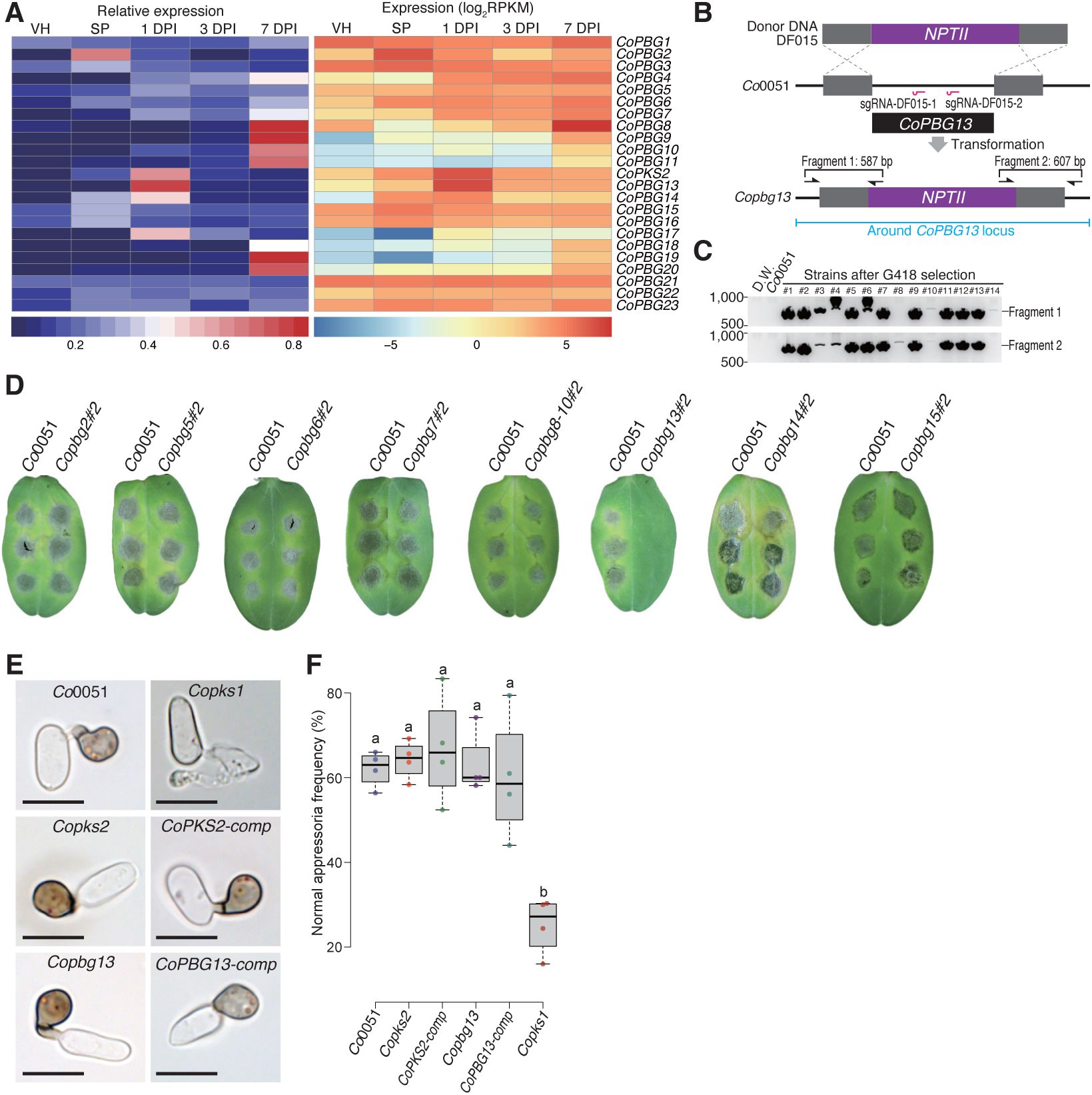
Characterization of genes within the *CoPKS2*-BGC. (**A**) Expression analysis of 23 genes in *CoPKS2*-BGC (BGC4). VH (vegetative hyphae) and SP (spore) were non-infection controls. Details of the heatmaps and samples are the same as those in Fig. 1A. (**B**) Schematics of *Copbg13* knockout. PCR-amplified DNA fragments were used as donor DNA. *NPTII* is a selection marker conferring G418/geneticin resistance. Grey boxes, magenta lines, and arrows indicate homology arms, sequences targeted by gRNAs of CRISPR-Cas9, and primers respectively. Details of the donor DNA, primers, and each Fragment are listed in Data S3, S4, and S5, respectively. (**C**) PCR screening of *Copbg13* mutants. The primer sets used are depicted in (B). Strains that exhibited both bands corresponding to Fragment 1 and Fragment 2 were selected as *Copbg13*. (**D**) Disease symptoms of *Co* mutant strains on cucumber. *Codmat4, Conrpsl9,* and *Conrps8* showed similar disease symptoms to *Co*0051, which served as a control. The same experiments depicted in Fig. 1B were performed. Three independent experiments yielded similar results. (**E**) Morphology of in vitro formed appressoria of each *Co* genotype. Scale bars represent 10 µm. (**F**) Frequency of normal appressoria of each *Co* genotype. Normal appressoria frequency was calculated as the percentage of normal appressoria in all counted spores. Normal appressoria were represented as that of *Co*0051 shown in (E). Each data point represents the normal appressoria frequency calculated from at least 50 appressoria. Four data points were used for each genotype (n=4). Different characters indicate a statistically significant difference with a p-value of less than 0.05 (Tukey HSD).

**Fig. S3.**
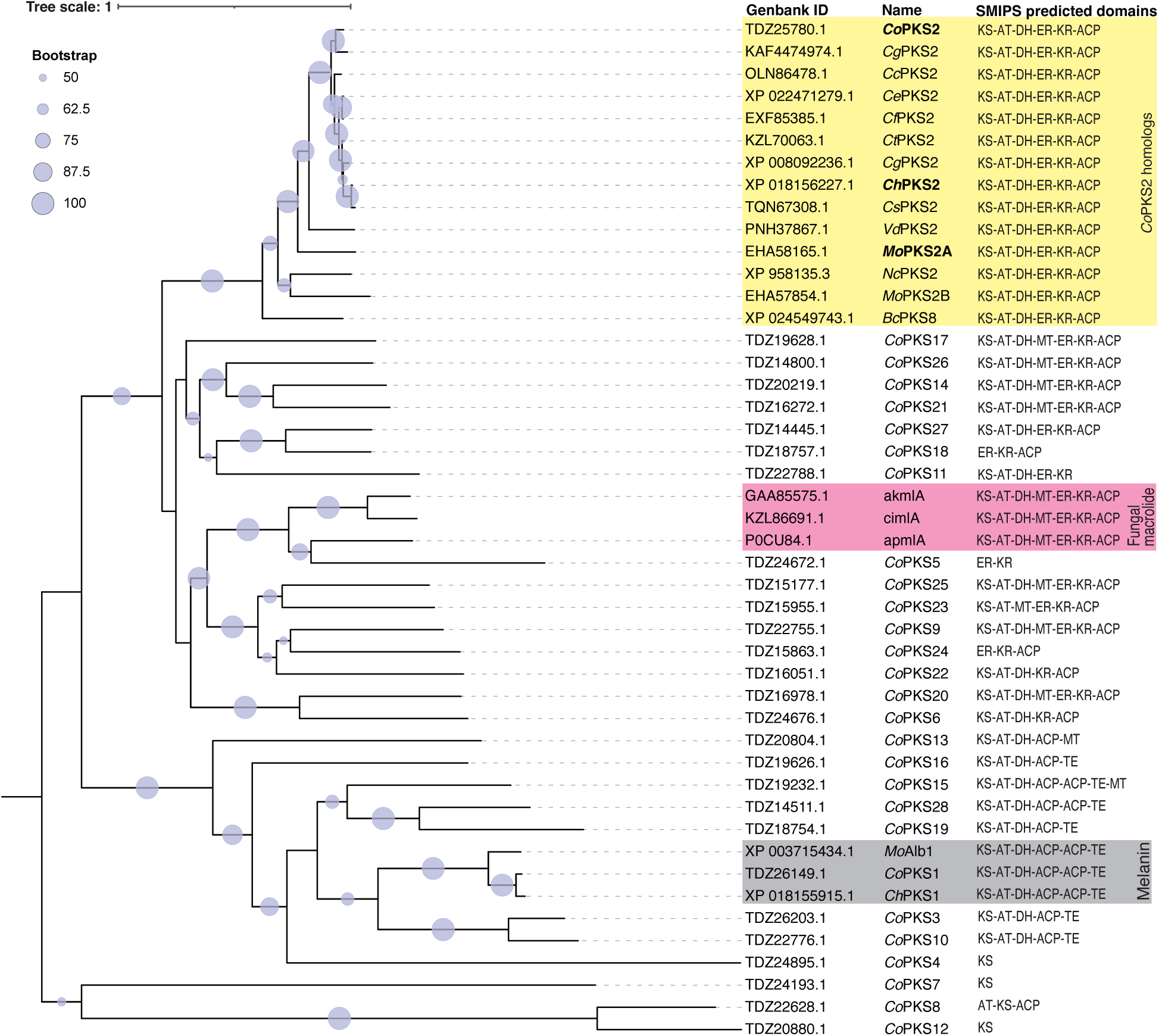
Phylogenetic analysis of PKS2 homologs and all *Co*PKSs. A phylogenetic tree of 46 PKSs was generated using maximum likelihood phylogenetic analysis. The breakdown of the 46 PKSs is as follows: 28 from *Co*; 3 from fungal macrolide-related PKSs (*62*, *63*); 2 from *Ch* and *Mo* involved in melanin synthesis (*27*, *64*); 13 PKS2 homologs. Circle sizes on branches represent percent support values out of 1,000 bootstrap replicates. Only bootstrap values greater than 50% support are shown as circles. Predicted domains in each PKS were identified using the SMIPS (*51*) software. The domains include KS, ketosynthase; AT, acyltransferase; DH, dehydratase; ER, enoylreductase; KR, ketoreductase; ACP, acyl carrier protein; MT, methyltransferase; TE, thioesterase. The first two letters of each protein name represent the following species: *Co*, *Colletotrichum orbiculare*, *Ce*, *Colletotrichum gloeosporioides*, *Cc*, *Colletotrichum chlorophyti*; *Cr*, *Colletotrichum orchidophilum*; *Cf*, *Colletotrichum fioriniae*; *Ct*, *Colletotrichum tofieldiae*; *Cg*, *Colletotrichum graminicola*; *Ch*, *Colletotrichum higginsianum*; *Cs*, *Colletotrichum shisoi*; *Vd*, *Verticillium dahliae*; *Nc*, *Neurospora crassa*; *Mo*, *Magnaporthe oryzae*; *Bc*, *Botrytis cinerea*; ak, *Aspergillus kawachii*; ci, *Colletotrichum incanum*; ap, *Arthrinium phaeospermum*.

**Fig. S4.**
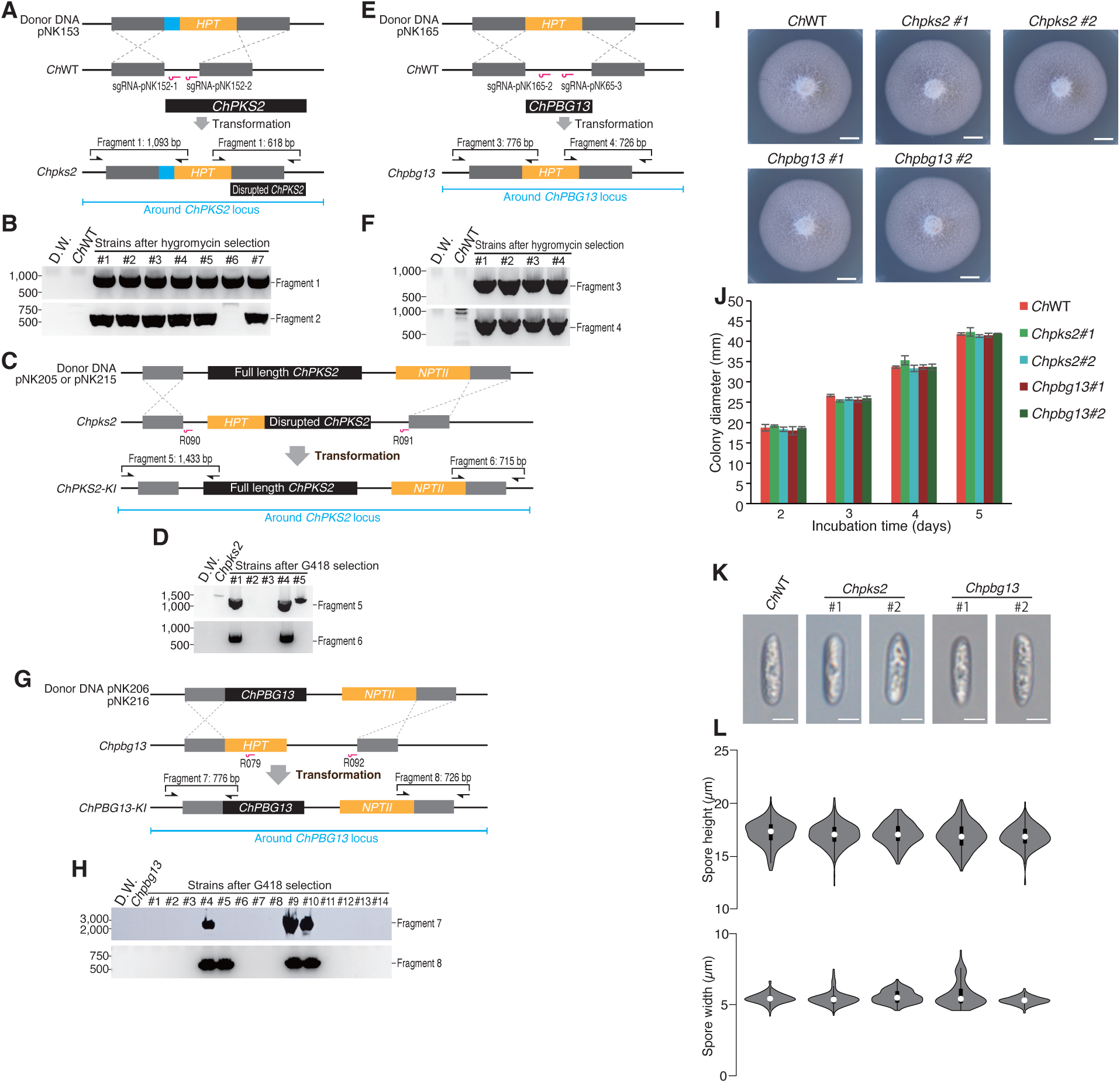
*ChPKS2* and *ChPBG13* knockouts do not affect colony and spore morphology. (**A, E**) Schematics of *ChPKS2* and *ChPBG13* knockout. *HPT* confers hygromycin resistance. Grey boxes represent homology arms, and magenta lines indicate CRISPR-Cas9 target sequences. (**B, F**) PCR screening of *Chpks2* or *Chpbg13* mutants. Primer sets from (A) or (E) were used. Strains showing bands corresponding to both Fragment 1 and Fragment 2, or to both Fragment 3 and Fragment 4, were selected as *Chpks2* or *Chpbg13* mutants, respectively. (**C, G**) Schematics of *ChPKS2* or *ChPBG13* knock-in. *NPTII* confers G418 resistance. Grey boxes indicate homology arms and magenta lines represent CRISPR-Cas9 target sequences. The 20 bp target sequence of gRNAs R090, R091, and R092 were excised from donor DNAs to prevent cleavage. (**D, H**) PCR screening of *ChPKS2-knockin (KI)* or *ChPBG13-KI* strains. Primer sets from (C) or (G) were used. Strains showing bands corresponding to both Fragment 5 and Fragment 6, or both Fragment 7 and Fragment 8, were selected as *ChPKS2-KI* or *ChPBG13-KI* strains, respectively. In A-H, details of the donor DNA, primers, and each Fragment are listed in Data S3, S4, and S5, respectively. (**I, J**) Colony morphology (I) and diameter (J) of each *Ch* strain on Mathur’s media. Error bars indicate ±SE (n=3). Scale bars: 5 mm. (**K, L**) Spore morphology (K), height, and width (H) of each *Ch* strain. Scale bars: 5 µm. Violin plots were based on measurements of at least 228 spores for each strain.

**Fig. S5.**
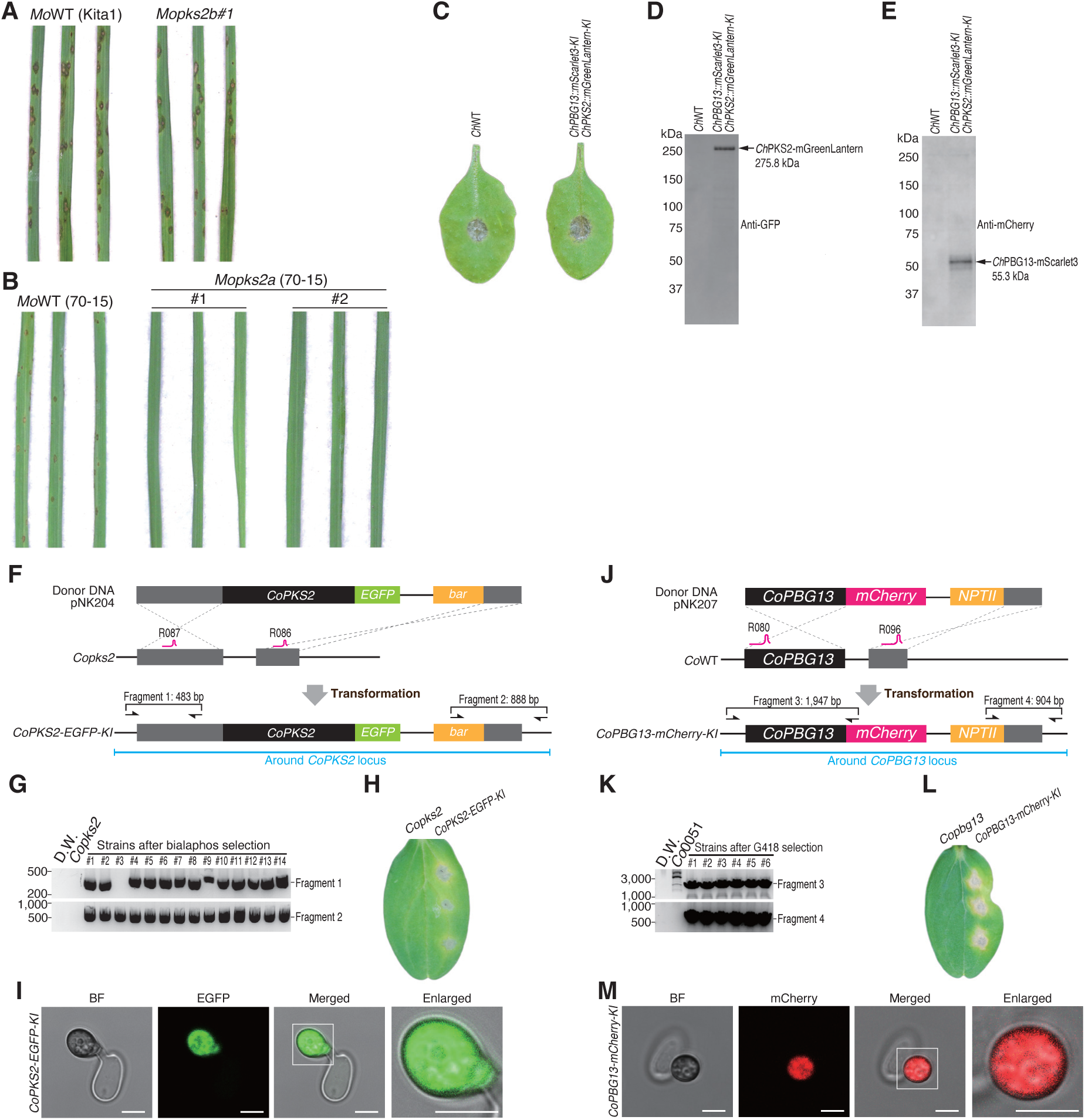
PKS2 and PBG13 localized to appressorial cytosol. (**A**) *Mopks2b* exhibited disease symptoms similar to *Mo*WT on rice. (**B**) *Mopks2a* (70-15 strain) showed no disease symptoms on rice. (**C**) Double knock-in, *ChPKS2-mGreenLantern-KI ChPBG13-mScarlet3-KI*, showed similar disease symptoms to *Ch*WT. *Ch* strains were inoculated onto *A. thaliana* Col-0, resulting in anthracnose symptoms. Six leaves were used for each *Ch* genotype (n=6). (**D, E)** Immunoblot showing detection of *Ch*PKS2-mGreenLantern and *Ch*PBG13-mScarlet3 fused proteins in samples isolated from in vitro induced appressoria of *Ch* strains. Signals were detected using an anti-GFP antibody (D) and an anti-mCherry antibody (E). (**F, J**) Schematics illustrating knock-in of *CoPKS2-EGFP* or *CoPBG13-mCherry*. Bialaphos and G418 resistance conferred by *bar* and *NPTII*, respectively. Grey boxes depict homology arms, and magenta lines indicate CRISPR-Cas9 target sequences. A black box corresponding to *CoPBG13* was used as a homology arm in (J). Details of the donor DNA, primers, and each Fragment are listed in Data S3, S4, and S5, respectively. (**G, K**) PCR screening of *CoPKS2-EGFP-knockin* (*-KI*) or *CoPBG13-mCherry-KI* strains using primer sets from (F) or (J). Strains showing bands for both Fragment 1 and Fragment 2 or both Fragment 3 and Fragment 4, were selected as *CoPKS2-EGFP-KI* and *CoPBG13-mCherry-KI*, respectively. (**H, I**) Disease symptom on cucumber leaves caused by *CoPKS2-EGFP-KI* and *CoPBG13-mCherry-KI.* Four cotyledons were used for each *Co* genotype (n=4). (**I, M**) Localization of *Co*PKS2-EGFP and *Co*PBG13-mCherry to appressorial cytosol. Images are single optical sections. Scale bars represent 5 µm. In (C, H, I), three independent experiments yielded similar results.

**Fig. S6.**
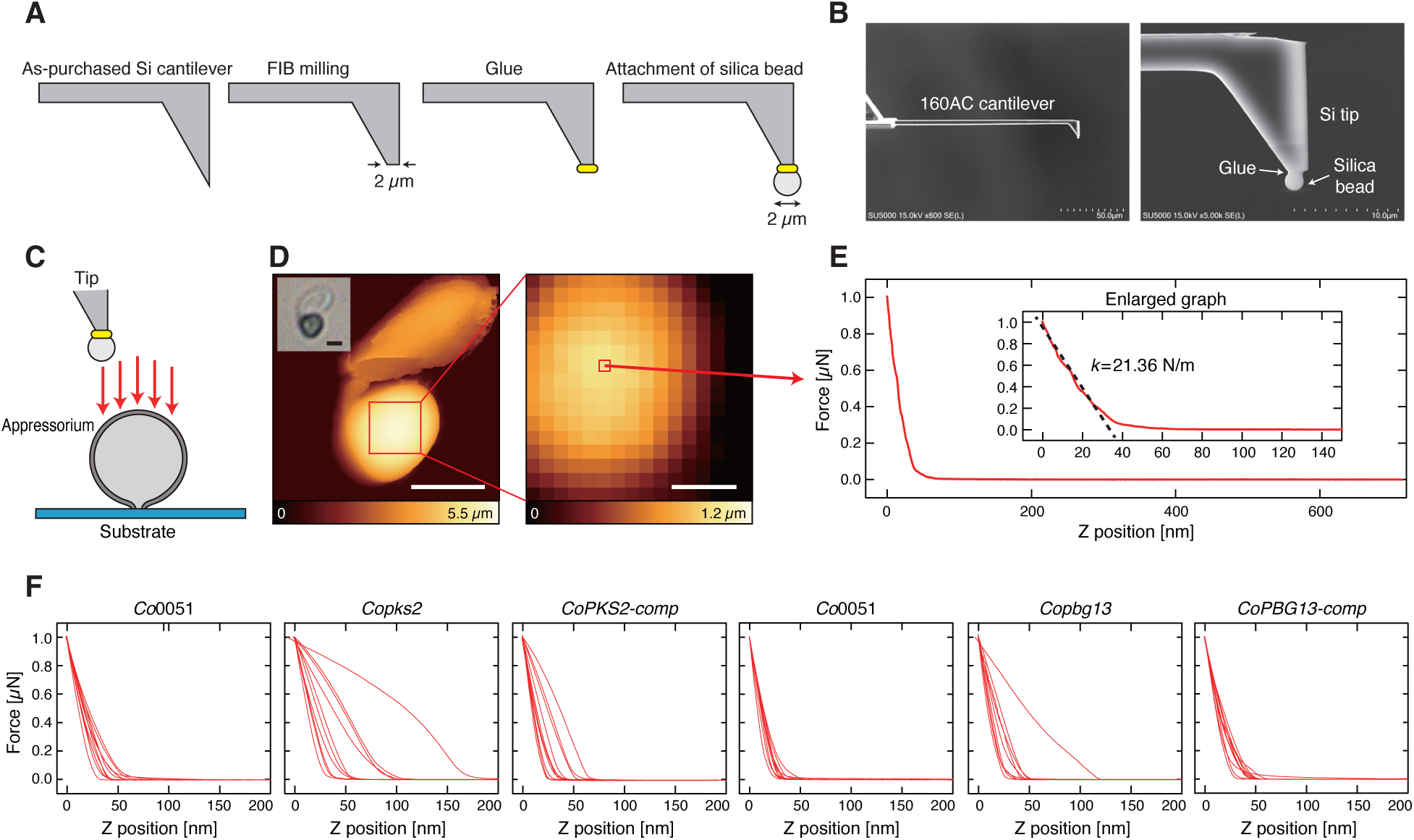
Stiffness measurement of appressorium by AFM. (**A**) Fabrication process of an AFM tip for stiffness measurements. (**B**) Scanning electron microscopy image of the fabricated tip. (**C**) Schematic illustration of AFM measurements. (**D**) Height images (inset: bright field optical microscopy image, scale bar: 5 μm) of an appressorium measured by AFM. (**E**) Force curve obtained at a central position on the upper surface of an appressorium (inset: enlarged graph with linear fitting for calculating the stiffness) using AFM. (**F**) The measured force curves (n=10) for appressoria of each genotype using AFM.

**Fig. S7.**
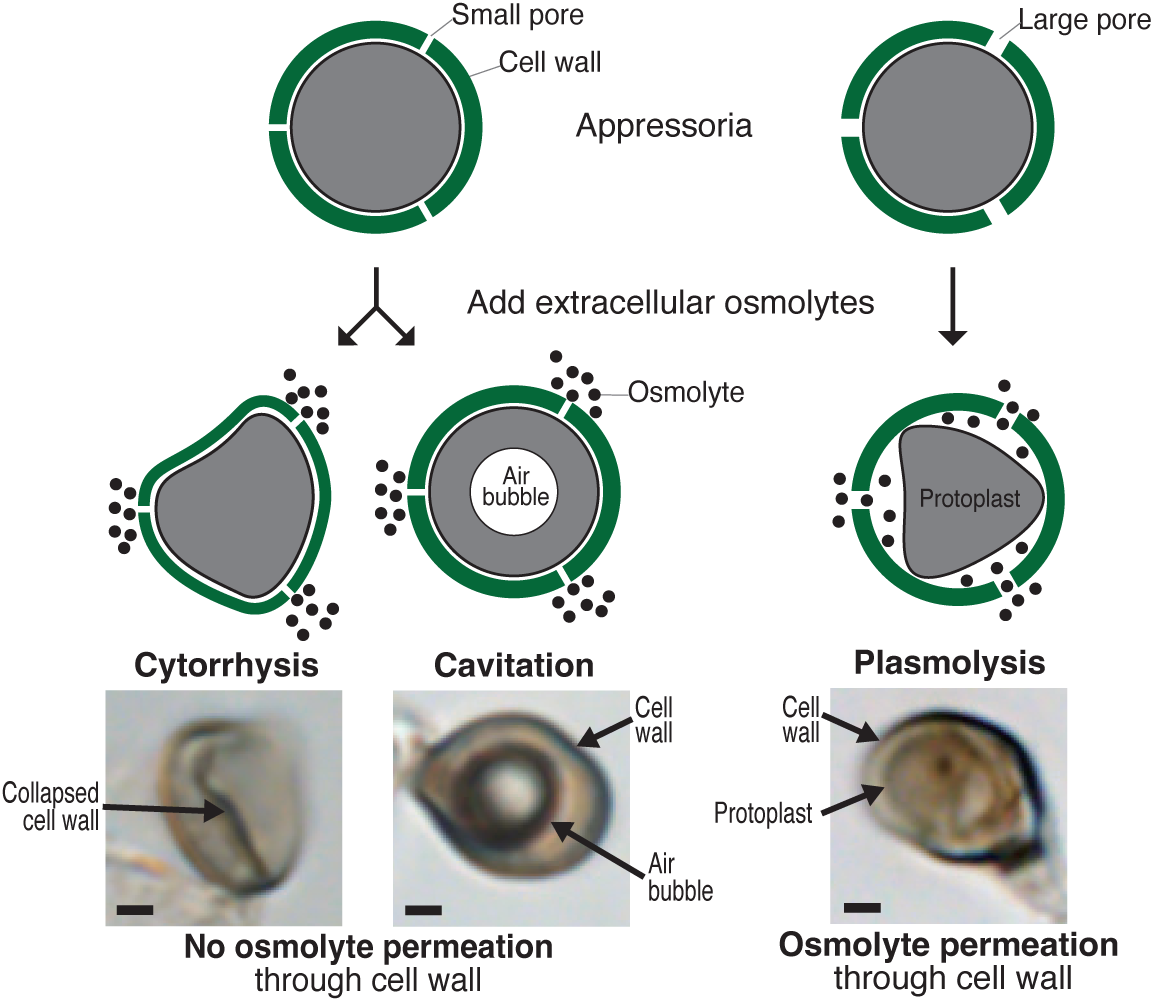
Interpretation of incipient cytorrhysis assay. Cytorrhysis or cavitation after treatment with the osmolyte (PEG-6000) indicates that the osmolyte cannot permeate through the cell wall because the pores are too small, while plasmolysis indicates that the cell wall pores are large enough to allow permeation the osmolyte. In the case of cytorrhysis, the cell wall collapses upon dehydration of the cell by osmosis, while in the case of cavitation the cell wall is too rigid to collapse, and dehydration causes formation of an air bubble in the cytoplasm. Representative microscope images of cytorrhysis, cavitation, and plasmolysis in *Co* appressoria. Scale bars represent 1 µm.

**Fig. S8.**
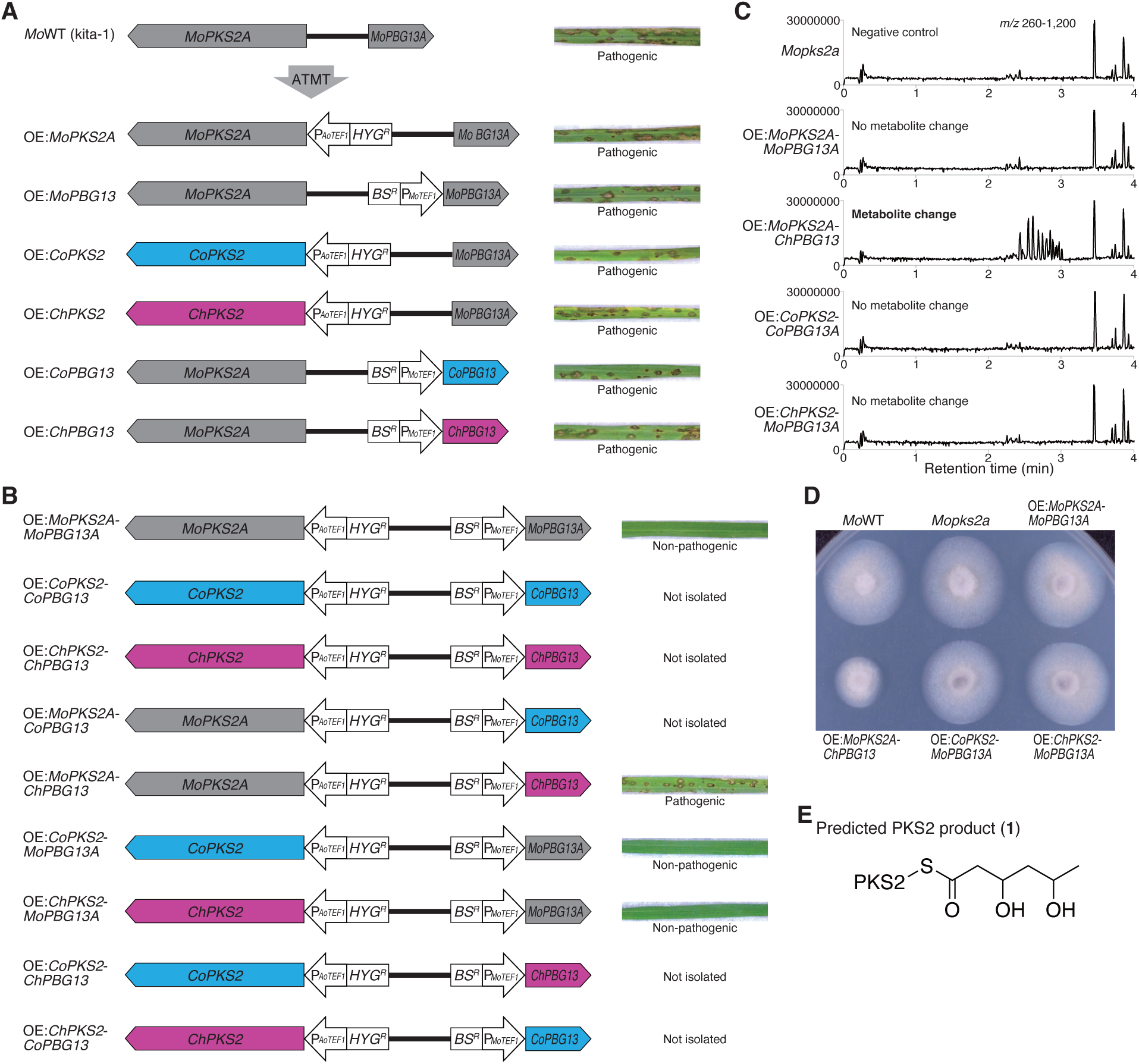
Homologous and heterologous expression of PKS2-PBG13 in *Mo*. (**A, B**) Construction and analysis of single (A) and double (B) overexpression strains of *Mo*. Infected rice leaves of each strain are shown. ATMT, *Agrobacterium tumefaciens*-mediated transformation; P*_AoTEF1_*, *Aspergillus oryzae TEF1* promoter; P*_MoTEF1_*, *Mo TEF1* promoter; *HYG^R^*, hygromycin B-resistance gene expression cassette; *BS^R^*, blasticidin S-resistance gene expression cassette. Details of each *Mo* strain are described in Data S6. (**C**) Metabolite analysis of double overexpression strains. *Mo* strains were cultured in YG medium for 7 d. Metabolites were analyzed by UPLC and detected by MS at *m/z* 260-1200. (**D**) *MoPKS2A-ChPBG13* overexpression strains showed slower growth. (**E**) Predicted PKS2 product (**1**).

**Fig. S9.**
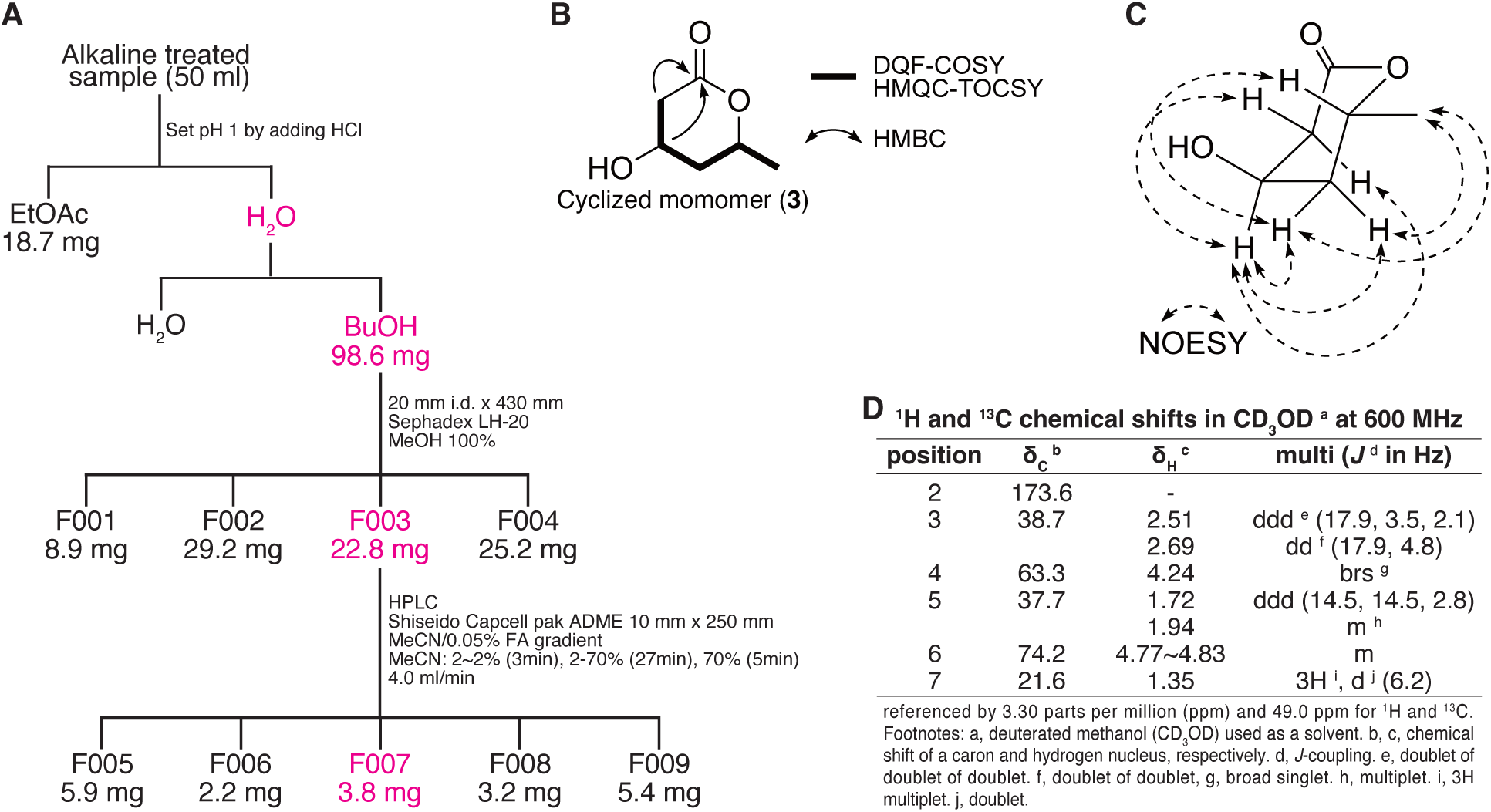
NMR analysis of cyclized monomer. (**3**) (**A**) The procedure for isolating cyclized monomer (**3**). (**B**) Selected 2D NMR correlations to identify the planar structure of **3**. (**C**) Selected NOESY correlations to confirm the relative stereochemistry. (**D**) ^1^H and ^13^C chemical shifts in CD_3_OD at 600 MHz.

**Fig. S10.**
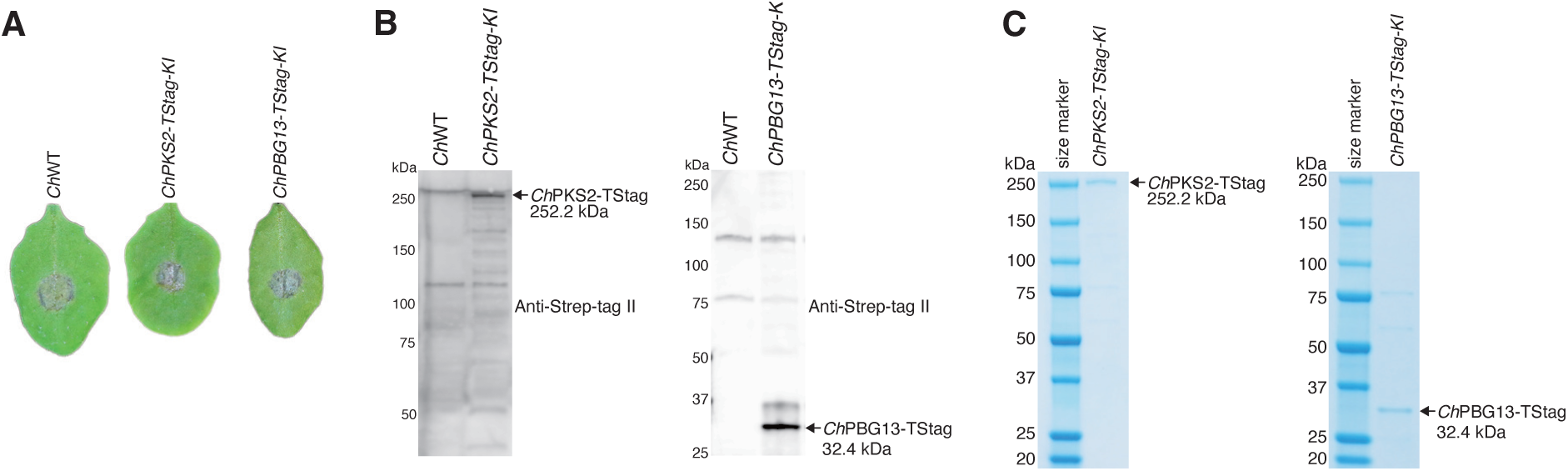
Purification of *Ch*PKS2-TStag and *Ch*PBG13-TStag proteins. (**A**) *ChPKS2-TStag-KI and ChPBG13-TStag-KI* strains exhibited disease symptoms on *A. thaliana* Col-0. Three leaves were used for each genotype (n=3). Three independent experiments yielded similar results. (**B**) Detection of *Ch*PKS2-TStag or *Ch*PBG13-TStag proteins from appressoria by immunoblotting. Anti-Strep-tag II antibody was used for detection. (**C**) Purified *Ch*PKS2-TStag or *Ch*PBG13-TStag proteins. Purified proteins were separated by SDS-PAGE and stained with Coomassie Blue.

**Fig. S11.**
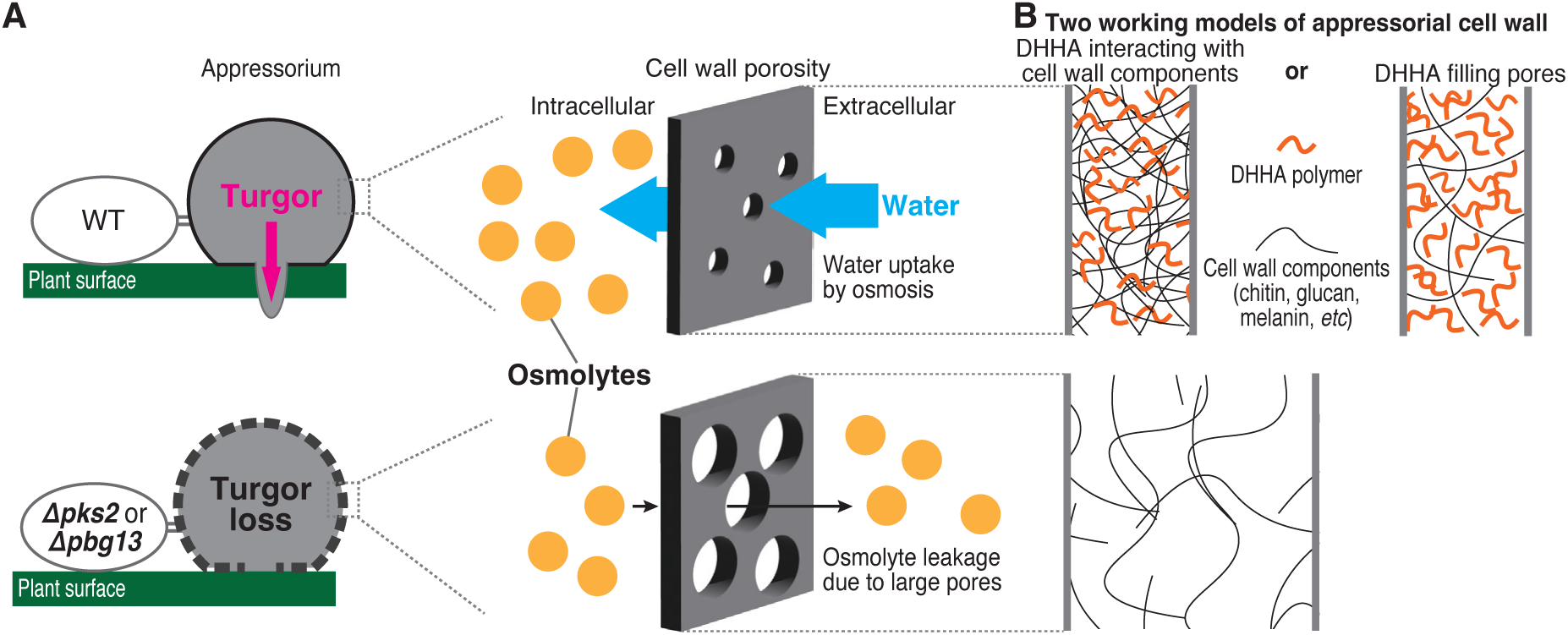
Schematic representation of turgor regulation by PKS2 and PBG13. **(A)** Schematic representation of turgor regulation by *PKS2* and *PBG13*. In a WT appressorium, the cell wall porosity is small enough to restrict osmolyte leakage, allowing water uptake by osmosis to generate high turgor pressure. In contrast, in the *pks2* or *pbg13* mutants, the cell wall porosity is larger due to the lack of DHHA polymers, allowing osmolytes to leak out of the cell, thereby preventing the generation of high turgor pressure. (**B**) Two working models of cell wall porosity regulation by DHHA. In the first model, DHHA polymers interact with cell wall components, such as polysaccharides (chitin and glucans) and DHN-melanin, making them more densely packed and thereby reducing wall porosity. In the second model, DHHA polymerizes independently to fill the spaces between these cell wall components, reducing the pore size.

**Table S1.**
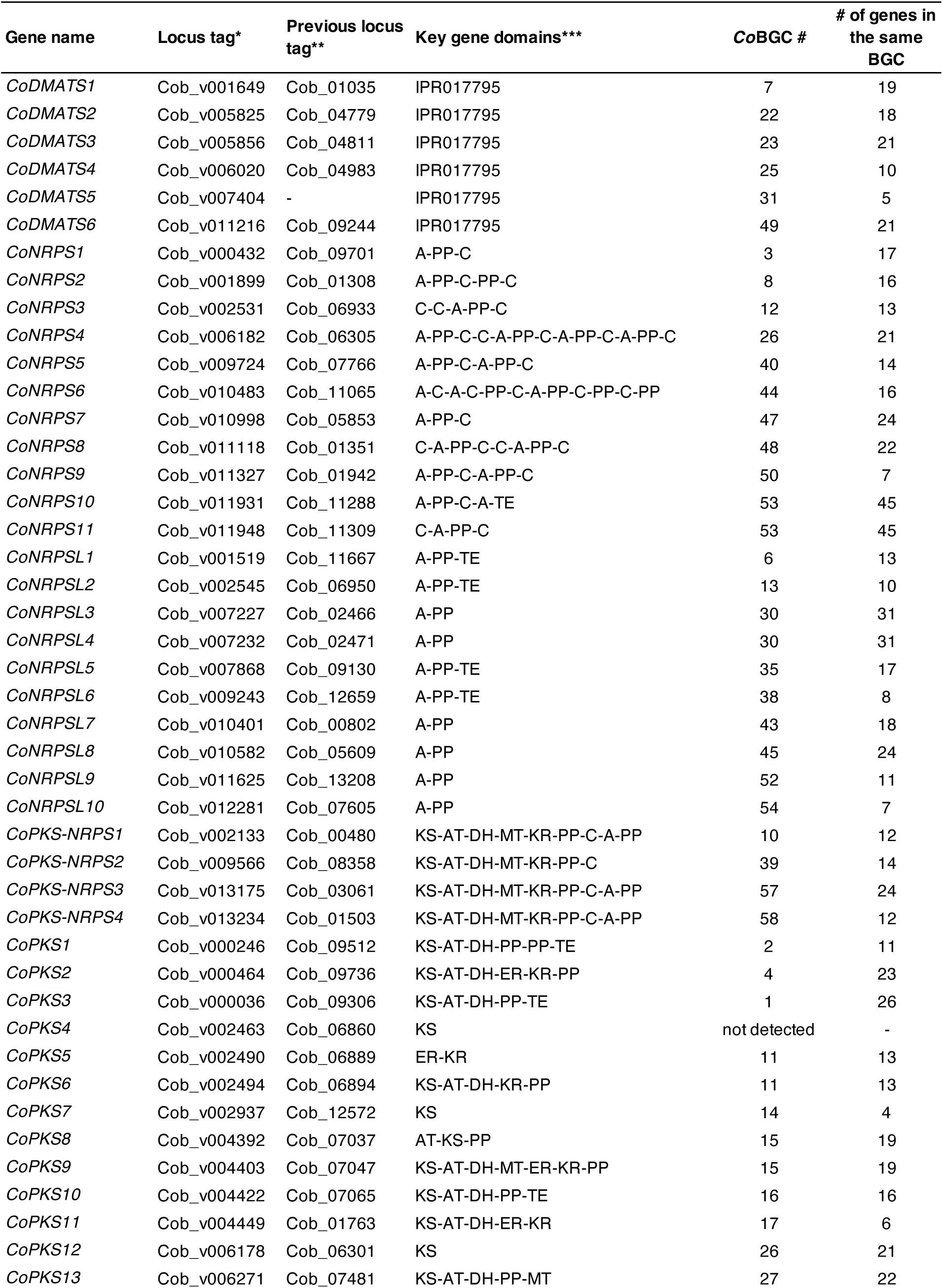
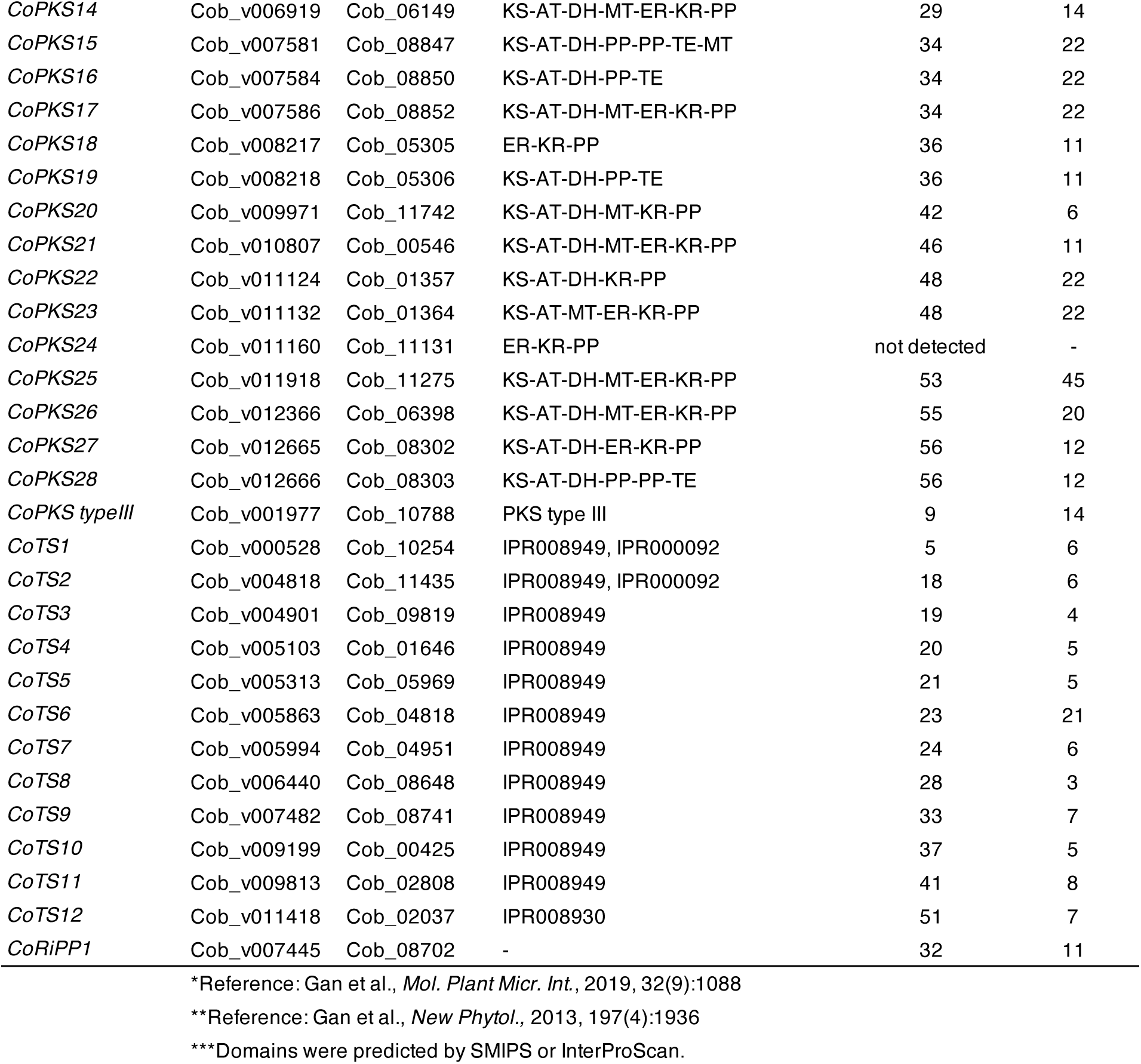
List of *Co*SMKGs.

**Table S2.**
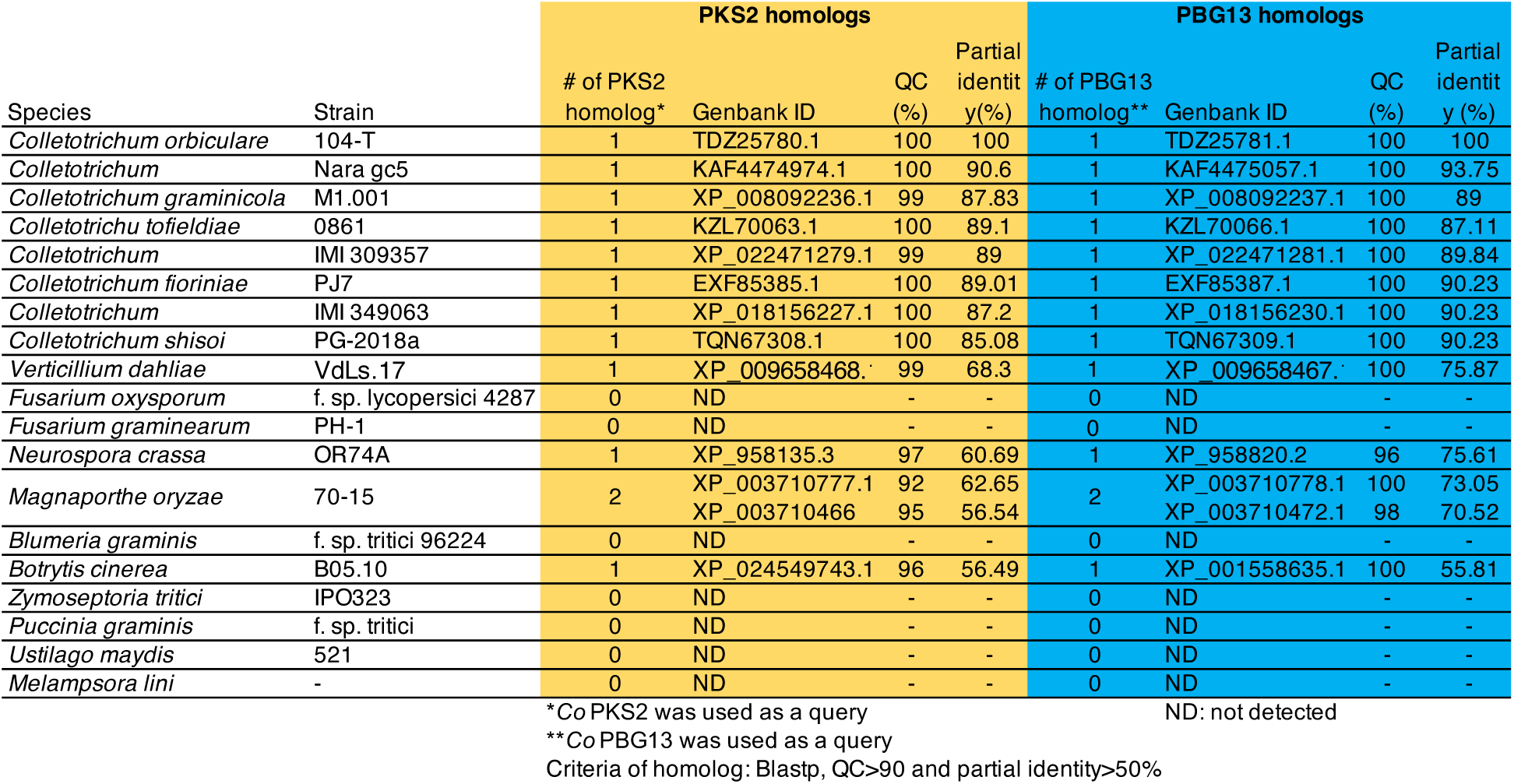
List of PKS2 and PBG13 homologs.

**Table S3.**
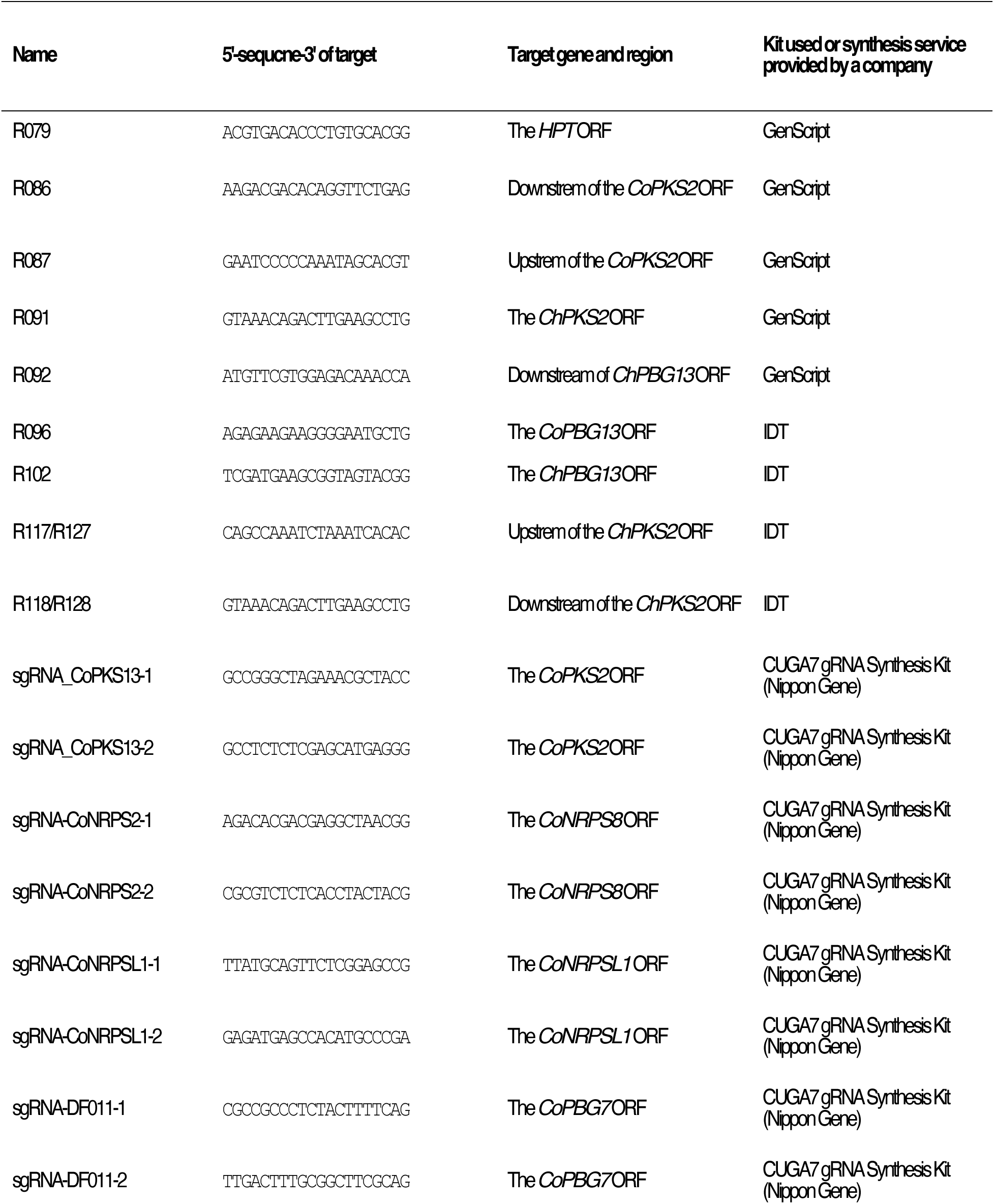
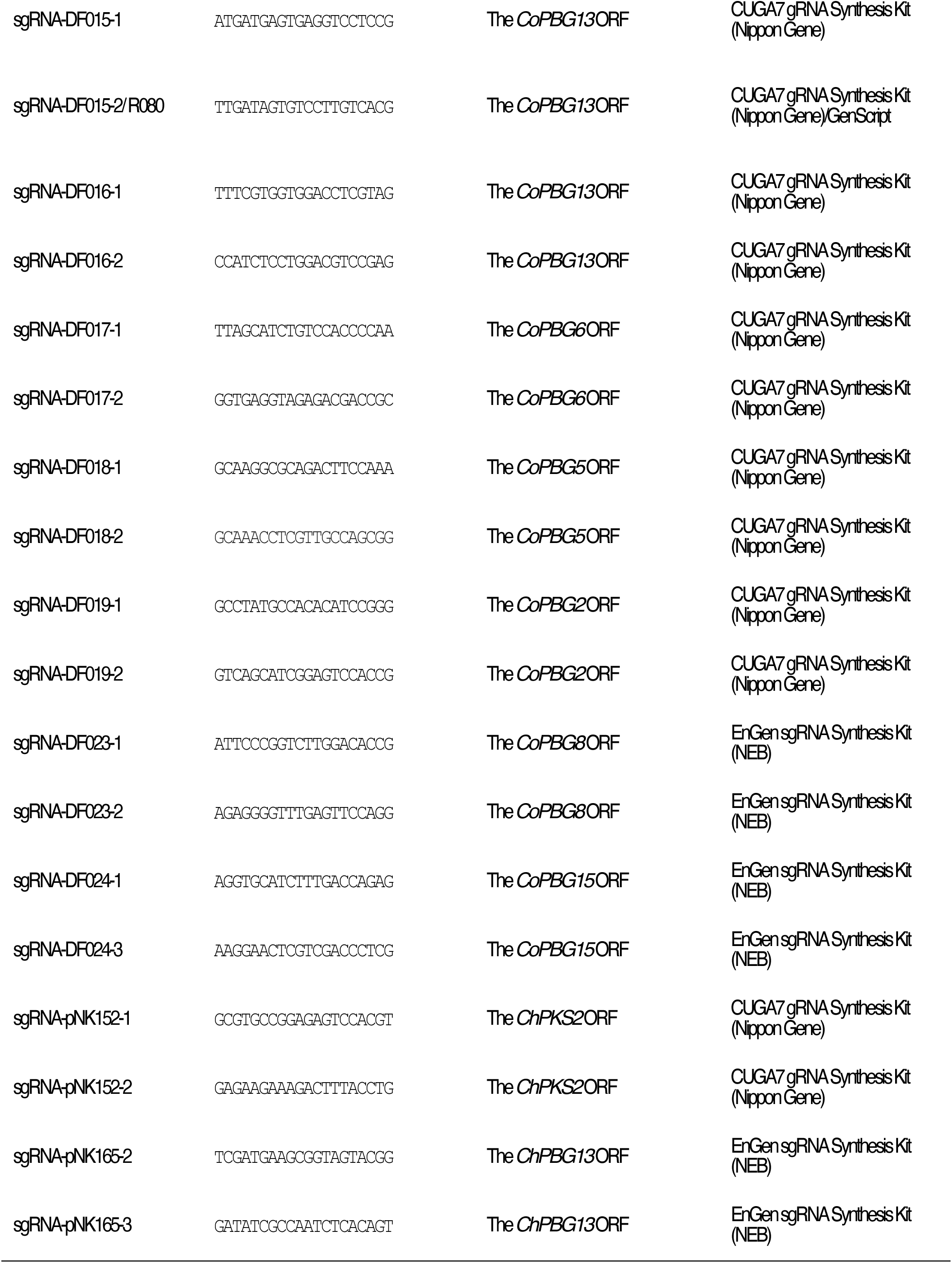
gRNA list.

**Table S4.**
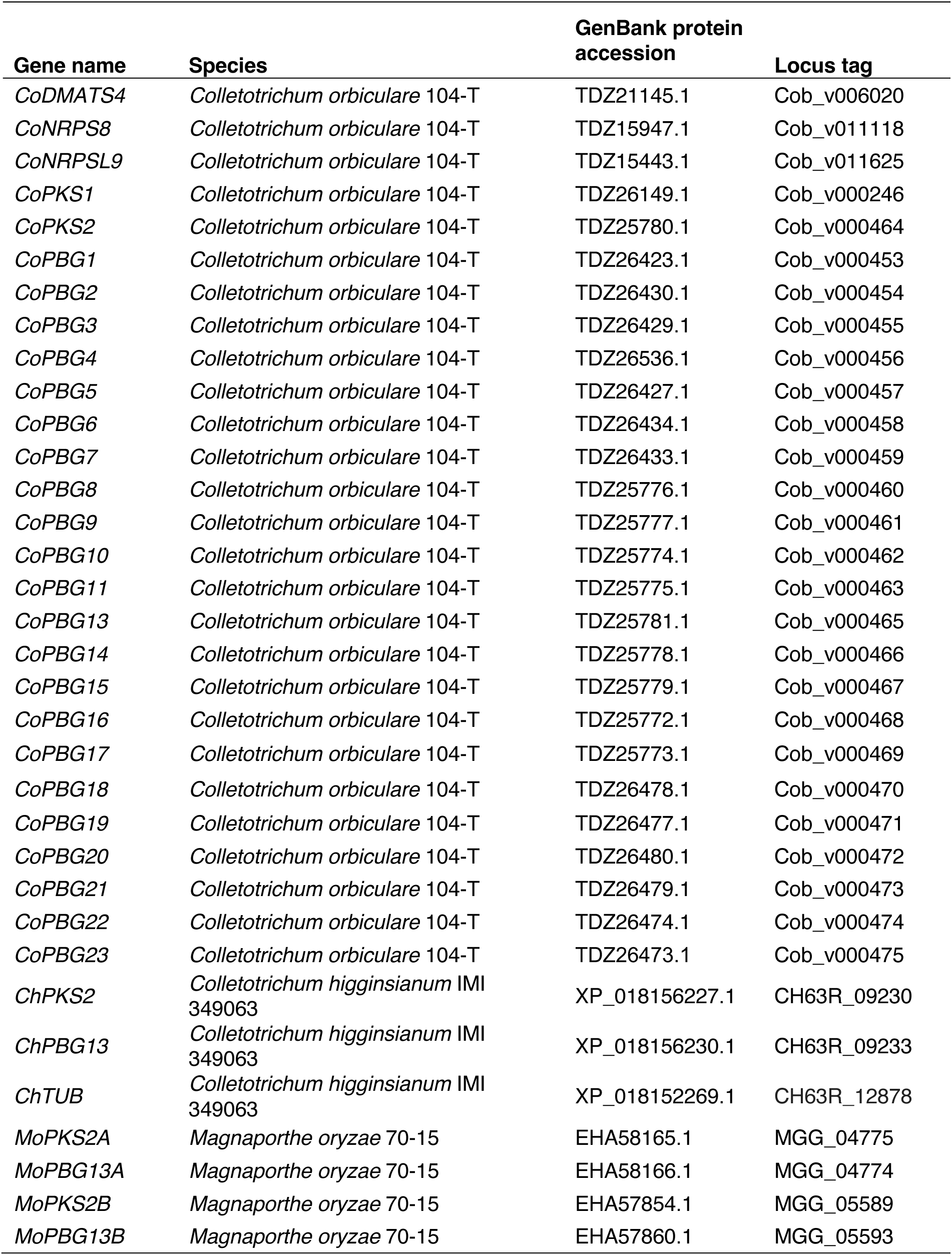
Accession numbers.

**Data S1. (separate file)**

List of *Co*BGCs

**Data S2. (separate file)**

RPKM and CPM of each *Co* gene

**Data S3. (separate file)**

DNA list

**Data S4. (separate file)**

Oligonucleotide list

**Data S5. (separate file)**

*Colletotrichum* strain list

**Data S6. (separate file)**

*Mo* strain list

